# Early transcriptional responses of human nasal epithelial cells to infection with Influenza A and SARS-CoV-2 virus differ and are influenced by physiological temperature

**DOI:** 10.1101/2023.03.07.531609

**Authors:** Jessica D. Resnick, Michael A. Beer, Andrew Pekosz

## Abstract

Influenza A (IAV) and SARS-CoV-2 (SCV2) viruses represent an ongoing threat to public health. Both viruses target the respiratory tract, which consists of a gradient of cell types, receptor expression, and temperature. Environmental temperature has been an understudied contributor to infection susceptibility and understanding its impact on host responses to infection could help uncover new insights into severe disease risk factors. As the nasal passageways are the initial site of respiratory virus infection, in this study we investigated the effect of temperature on host responses in human nasal epithelial cells (hNECs) utilizing IAV and SCV2 *in vitro* infection models. We demonstrate that temperature affects SCV2, but not IAV, viral replicative fitness and that SCV2 infected cultures are slower to mount an infection-induced response, likely due to suppression by the virus. Additionally, we show that that temperature not only changes the basal transcriptomic landscape of epithelial cells, but that it also impacts the response to infection. The induction of interferon and other innate immune responses were not drastically affected by temperature, suggesting that while the baseline antiviral response at different temperatures remains consistent, there may be metabolic or signaling changes that affect how well the cultures are able to adapt to new pressures such as infection. Finally, we show that hNECs respond differently to IAV and SCV2 infection in ways that give insight into how the virus is able to manipulate the cell to allow for replication and release. Taken together, these data give new insight into the innate immune response to respiratory infections and can assist in identifying new treatment strategies for respiratory infections.

## INTRODUCTION

Two of the most severe pandemics in recent history-1918 Influenza and COVID-19-were caused by respiratory pathogens ^1^. Influenza A virus (IAV) is an 8-segmented, negative sense RNA virus belonging to the *Orthomyxoviridae* family that continues to cause seasonal epidemics. On average, 8.3% of the United States population experiences influenza infection each year, and the 2022-2023 season is already shaping up to be one of the most severe in recent history^2,3^. Recent studies have tried to elucidate what factors may predispose some individuals to have such severe reactions to IAV infection. These studies have not only included investigations into components of the virus and adaptive immunity of the host, but also age, sex, microbiome, lifestyle and genetic variation^4^.

SARS-CoV-2 (SCV2) virus is a non-segmented, positive sense RNA virus belonging to the *Coronaviridae* family and the causative agent of COVID19 disease^5,6,7,8,9^. As of December 12, 2022, the Johns Hopkins dashboard reports 649,207,625 total cases and 6,653,264 total deaths worldwide, although this is likely an underestimation due to incomplete reporting ^9^. While the current fatality rate for COVID19 continues to be unclear due to virulence of different variants and the impact of pre-existing immunity from vaccination, early estimates ranged from 0.4-3.6% compared to 0.1% for influenza^10^. However, the continued burden on intensive care units suggests instances of severe disease are more wide-spread. While IAV infection can lead to severe disease phenotypes such as pneumonia and acute respiratory distress syndrome (ARDS) as well as diarrhea and abdominal pain (usually in children), SCV2 infection additionally can lead to severe endothelial damage, vascular thrombosis with microangiopathies, occlusion of vascular capillaries, and cytokine storm characterized by high levels of IL-6 ^11^. Studies have also shown that risk factors for severe COVID19 are similar but distinct to those for Influenza, and include factors such as being male, advanced age, obesity, genetic risk factors, and autoantibodies ^32,33,34^.

The first barrier to respiratory infection lies in the innate immune responses of the respiratory epithelium. However, to date most studies have used blood samples from patients in order to evaluate infection severity and disease correlates ^14,15^. While some studies have addressed the respiratory epithelial cell responses to virus infection, few studies have accounted for the physiological differences in epithelial cell types and temperature present between the upper and lower respiratory tract that could contribute to differences in the severity of infection, with a majority of SCV2 studies being done in bronchial epithelial cell cultures ^61,62,63^. Additionally, airway epithelial cell types are significantly more diverse than previously believed, and a recent single-cell analysis have shown that different cell types can have more of an immune regulatory profile, leading to new questions about cell specific functions in the innate immunity of the airway epithelium^19^.

IAV primarily targets airway and alveolar epithelial cells that express sialic acid receptors ^16^. However, individual viruses have slightly different receptor preferences. The clearest example of this is avian influenza virus preference for an alpha 2,3 linked sialic acid, which is more common in avian intestinal tracts, compared to human influenza viruses that prefer alpha 2,6 linkages. But most viruses are not this dichotomous, and the ability to utilize different glycans lies on a spectrum^17^. These glycans are most often expressed on ciliated cells, and although other epithelial cell types are susceptible to infection, a majority of ciliated cells will become infected with IAV ^18^. In the lower respiratory tract, which contains fewer ciliated cells relative to the upper respiratory tract, IAV has been found to bind predominately to type 1 pneumocytes in addition to the few ciliated cells that are present ^18^.

In contrast, the SARS-CoV2 virus utilizes the ACE2 receptor, along with proteolytic priming by proteases such as TMPRSS2, to enter target cells^22^. Single nuclei analysis of lung and bronchial cells revealed that these two proteins are most often co expressed in AT2 cells in lung tissue and secretory 3 cells, an intermediate of club and goblet cells, in hBEC cultures^23^. However, more recent studies have shown that expression of entry receptors and co-mediators are poor predictors of susceptibility ^24^. Long term infection studies utilizing hBEC cultures suggest the major targets of SARS-CoV2 virus are ciliated and goblet cells, while basal and club cells are not permissive to infection^25^.

Response to IAV infection have been observed to be more cell-specific than virus-specific ^20^. For example, a study by Taye, et. al. showed that infection with both human and avian IAV strains elicited similar antiviral, pro-apoptotic and inflammatory signatures regardless of virus origin^20^. Severe disease is often associated with lower cellular expression of antiviral response genes, lower expression of IFN-related pathways, and overexpression of inflammatory pathways ^14^. This unregulated inflammation leads to lung injury, which has been shown to persist for up to 2 months in mice^21^. Additionally, multiple studies have shown that both SARS-CoV-2 and IAV infection of human airway epithelial cell cultures induce IP10 signaling and interferon responses, but that the overall inflammatory response is attenuated in SARS-CoV-2 infected cultures as compared to IAV infected cultures^26, 67^.

The structure of the human respiratory tract is a gradient of cell types, receptor expression, and temperature ^27,28,29^. The upper respiratory tract has a physiological temperature of 33°C, while the lower respiratory tract is maintained at core body temperature of 37°C. During infection and the resulting fever response, these temperatures can rise to as high as 39°C. Temperature has previously been shown to impact both viral replication and antiviral responses during rhinovirus infection, where higher induction of antiviral factors inhibits replication at 37°C in mice ^30^. However, other studies done in HeLa cells showed no such difference ^30,31^. Therefore, it is important to utilize physiologically relevant model systems along with clinical isolates of viruses to identify whether temperature may be an important modulator of severe respiratory disease.

As the nasal passageways are the initial site of respiratory virus infection, in this study we investigated the effect of temperature on host responses in primary, differentiated human nasal epithelial cell (hNECs) cultures utilizing IAV and SCV2 infection models using bulk RNAseq. We demonstrate that temperature affects viral replicative fitness as well as influences hNEC responses to infection. Additionally, we show that SCV2 infected cultures are slower to mount an infection-induced transcriptional response compared with IAV. These data indicate that physiological ranges of temperature should be accounted for when evaluating host responses to infection.

## METHODS

### Cell Culture

VeroE6TMPRSS2 cells (VT; RRID: CVCL_YQ49) were obtained from the cell repository of the National Institute of Infectious Diseases, Japan and are described ^41^. VT cells were cultured in Dulbecco’s Modified Eagle Medium (DMEM, Gibco) with 10%fetal bovine serum (FBS, Gibco Life Technologies), 100U penicillin/mL with 100ug streptomycin/mL (Quality Biological), 2mM L-Glutamine (Gibco Life Technologies), and 1mM Sodium Pyruvate (Sigma) at 37°C with air supplemented with 5% CO2. The infectious medium (IM) used in all SCV2 infections of VT cells consisted of DMEM with 2.5% FBS, 100U penicillin/mL with 100ug streptomycin/mL, 2mM L-Glutamine, and 1m M Sodium Pyruvate.

Madin-Darby canine kidney (MDCK) cells were cultured in Dulbecco’s Modified Eagle Medium (DMEM, Sigma-Aldrich) with 10% fetal bovine serum (FBS, Gibco Life Technologies), 100U penicillin/mL with 100 μg streptomycin/mL (Quality Biological), and 2 mM L-Glutamine (Gibco Life Technologies) at 37 °C with air supplemented with 5% CO2. The Infectious medium (IM) used in all IAV infections of MDCK cells consisted of DMEM with 4ug/mL N-acetyl trypsin (NAT), 100u/ml penicillin with 100ug/ml streptomycin, 2 mM L-Glutamine and 0.5% bovine serum albumin (BSA) (Sigma).

Human nasal epithelial cells (hNEC) (Promocell, lot 466Z004 and 453Z019) were grown to confluence in 24-well Falcon filter inserts (0.4-uM pore; 0.33cm2; Becton Dickinson) using PneumaCult™-Ex Plus Medium (Stemcell). Cultures derived from donor 453Z019, a 32 year old Caucasian male and donor 466Z004, a 43 year old Caucasian male, were used for initial growth curves and cytokine responses. Cultures derived from donor 466Z004 only were used for the RNA-sequencing experiment.. Confluence was determined by a transepithelial electrical resistance (TEER) reading above 250Ω by Ohm’s law method ^42^ and by examination using light microscopy and a 10x objective. The cells were then differentiated at an air-liquid interface (ALI) before infection, using ALI medium as basolateral medium as previously described^43,44^. Briefly, both apical and basolateral media were removed and ALI differentiation media (Stem Cell Technologies, Pneumacult ALI Basal Medium) supplemented with 1X ALI Maintenance Supplement (StemCell Technologies), 0.48 ug/mL Hydrocortisone solution (StemCell Technologies), and 4 ug/mL Heparin sodium salt in PBS (StemCell Technologies) was replaced on the basolateral side only. Fresh media was given every 48 hours. Once mucus was visible, apical washes were performed weekly with PBS to remove excess mucus. Cells were considered fully differentiated after 3 weeks and when cilia were visible using light microscopy and 10x objective. All cells were maintained at 37°C in a humidified incubator supplemented with 5% CO2.

### Virus Seed Stock and Working Stock Generation

The SARS-CoV-2 virus used in this study, designated SARS-CoV-2/USA/MDHP-8/2020 (B.1), was isolated from samples obtained through the Johns Hopkins Hospital network^64^. For virus working stocks, VT cells in a T75 or T150 flask were infected at an MOI of 0.001 with virus diluted in IM. After a one hour incubation at 33C, the inoculum was removed and IM added (10 ml for T75 and 20 ml for a T150 flask). When cytopathic effect was seen in approximately 75% of the cells, the supernatant was harvested, clarified by centrifugation at 400g for 10 minutes, aliquoted and stored at −65C.

The Influenza A Virus used was A/Baltimore/R0243/2018 (H3N2) (3C.3a) and was also isolated from samples obtained through the Johns Hopkins Hospital network as part of the CEIRS network ^65^. For virus working stocks, MDCK cells in a T150 flask were infected at an MOI of 0.001 with virus diluted in IM. After one hour, the inoculum was removed and fresh IM was added. When cytopathic effect was seen in approximately 50% of cells, the supernatant was harvested, aliquoted, and stored at −65°C.

### TCID_50_ Assay

VT or MDCK cells were grown to 90-100% confluence in 96-well plates. After being washed twice with PBS+, ten-fold serial dilutions of the viruses in IM were made and 20uL of each dilution was added to 6 wells. The plates were incubated at 37°C with 5% CO2 for 5 days. The cells were fixed by adding 75uL of 4% formaldehyde in PBS per well overnight and then stained with Napthol Blue Black solution overnight. Endpoint values were calculated by the Reed-Muench method^45^.

### Multiplicity of Infection (MOI) infections

For hNEC infections, an MOI of 0.1 TCID50 per cell was used. The basolateral media was collected, stored at −65°C, and replaced with fresh media every 48 hours. The apical side of the transwell was washed 3 times with correspond IAV or SCV2 IM (mock used SCV2-IM), with a 10 minute incubation at 37°C in between each. The virus inoculum was diluted in IM and 100uL was added to the apical side of cells and allowed to incubate for 2 hours. The inoculum was then removed, the cells washed 3 times with PBS-, and returned to the incubator. At 48 hours post infection, a 10 minute apical wash was performed with IM and collected and stored at −65°C. Infectious virus particle production in apical washes was quantified using TCID50 on VT or MDCK cells for SARS-CoV2 and Influenza A Viruses respectively.

### Cytokine Secretion

Secreted interferons, cytokines, and chemokines were quantified from the basolateral samples at 48 and 96 hours post infection from the hNEC infections. Measurements were performed using the V-Plex Human Chemokine Panel 1 (CCL2, CCL3, CCL4, CCL11, CCL17, CCL22, CCL26, CXCL10, and IL-8) (Meso Scale Discovery) and the DIY Human IFN Lambda 1/2/3 (IL-29/28A/28B) ELISA (PBL Assay Science) according to the manufacturer’s instructions. This panel has been used previously to characterize IAV infections of epithelial cells^43,68,69^. Each sample was analyzed in duplicate. Heatmaps were generated and hierarchical clustering was performed using the R package “pheatmap”^46^.

### RNA-Sequencing and Analysis

Total RNA was extracted and purified from hNECs using Trizol reagent and the PureLink RNA Mini kit, including on-column DNAse treatment (Invitrogen/ThermoFisher). Quantitation of Total RNA was performed with the Qubit BR RNA Assay kit and Qubit Flex Fluorometer (Invitrogen/ThermoFisher), and quality assessment performed by RNA ScreenTape analysis on an Agilent TapeStation 2200. Unique Dual-index Barcoded libraries for RNA-Seq were prepared from 100ng Total RNA using the Universal Plus Total RNA-Seq with NuQuant Library kit (Tecan Genomics), according to manufacturer’s recommended protocol. Library amplification was performed for 16 cycles, as optimized by qPCR. Quality of libraries was assessed by High Sensitivity DNA Lab Chips on an Agilent BioAnalyzer 2100. Quantitation was performed with NuQuant reagent, and confirmed by Qubit High Sensitivity DNA assay, on Qubit 4 and Qubit Flex Fluorometers (Invitrogen/ThermoFisher). Libraries were diluted, and equimolar pools prepared, according to manufacturer’s protocol for appropriate sequencer. An Illumina iSeq Sequencer with iSeq100 i1 reagent V2 300 cycle kit was used for final quality assessment of the library pool. For deep RNA sequencing, a 200 cycle (2×100bp) Illumina NovaSeq S2 run was performed at Johns Hopkins Genomics, Genetic Resources Core Facility, RRID:SCR_018669.

iSeq and NovaSeq data files were uploaded to the Partek Server and analysis with Partek Flow NGS software, with RNA Toolkit, was performed as follows: pre-alignment QA/QC and trimming of reads. Following this, sequences were uploaded to the Beer lab cluster for further analysis. Sequences were first checked for quality using FastQC^48^. All sequences were determined to be of good quality and were then aligned using HISAT2 to the GRCH38 genome^49^, ^59^. Sequences were also aligned to whole genome sequence of stock viruses obtained from the Influenza Research Database and NCBI^50.51^. SAM files were then converted to BAM using samtools^52^. A gene-count matrix was then generated from BAM files using “featureCounts” R package, and differential expression analysis was performed using “DESeq2” in R^53,54^. Pathway analysis of differentially expressed genes was also performed using the R packages “clusterProfiler”, “gProfiler”, and MSigDB^55,56,57^. Heatmaps were generated and hierarchical clustering was performed using the R package “pheatmap”^46^. Upset plot was generated using the R package “UpSetR”^47^. Other plots were made in base R or using the R package “ggplot”^58^. For detailed methods and a full list of packages used please see https://github.com/JRes9/Res-nicketal_IAVvSCV2temperature_2023..

All sequence files and sample information have been deposited at NCBI Sequence Read Archive, NCBI BioProject: PRJNA925547.

## RESULTS

### Temperature-dependent replication in IAV and SCV2 on human nasal epithelial cells

To determine if recent clinical isolates of IAV and early isolates of SCV2 show temperature-dependent replication, low MOI multistep growth curves were performed on human nasal epithelial cell (hNEC) cultures at either 33°C, a temperature consistent with the upper respiratory tract, or 37°C, a temperature consistent with the lower respiratory tract (Figure 1). While IAV showed no significant differences in replication kinetics due to temperature, SCV2 showed faster initial replication at 37°C compared to 33°C (Figure 1A and 1B). Additionally, SCV2 showed significantly slower replication kinetics compared to IAV at 33°C (Figure 1C) and 37°C (Figure 1D). Basolateral supernatant from mock, IAV, or SCV2 infected hNEC cultures was collected 48 and 96 HPI and secreted interferon, cytokines, and chemokines related to the proinflammatory response were measured by ELISA and MSD assay as has been previously described to evaluate epithelial responses to infection^43,68,69^. (Figure 2). While cytokine production and release is directional in polarized epithelial cells, previous research has shown that apical versus basolateral release differs mainly in quantity, rather than type, of cytokine released^43,68^. Values for each condition were averaged and scaled to calculate Z-score, and then hierarchical clustering was performed to identify patterns in the data. Overall, IAV infected hNECs secreted higher amounts of chemokines, cytokines, and interferon than SCV2 infected hNECs. Additionally, higher temperatures during infection and later timepoints correlated with higher production. Finally, only late, high temperature SCV2 infected hNECs clustered with any IAV infected samples. All other SCV2 infected samples, even with high infectious virus loads, clustered with mock infected samples suggesting a dampened or delayed innate immune response to SCV2 infection in epithelial cells.

**Figure 1:**
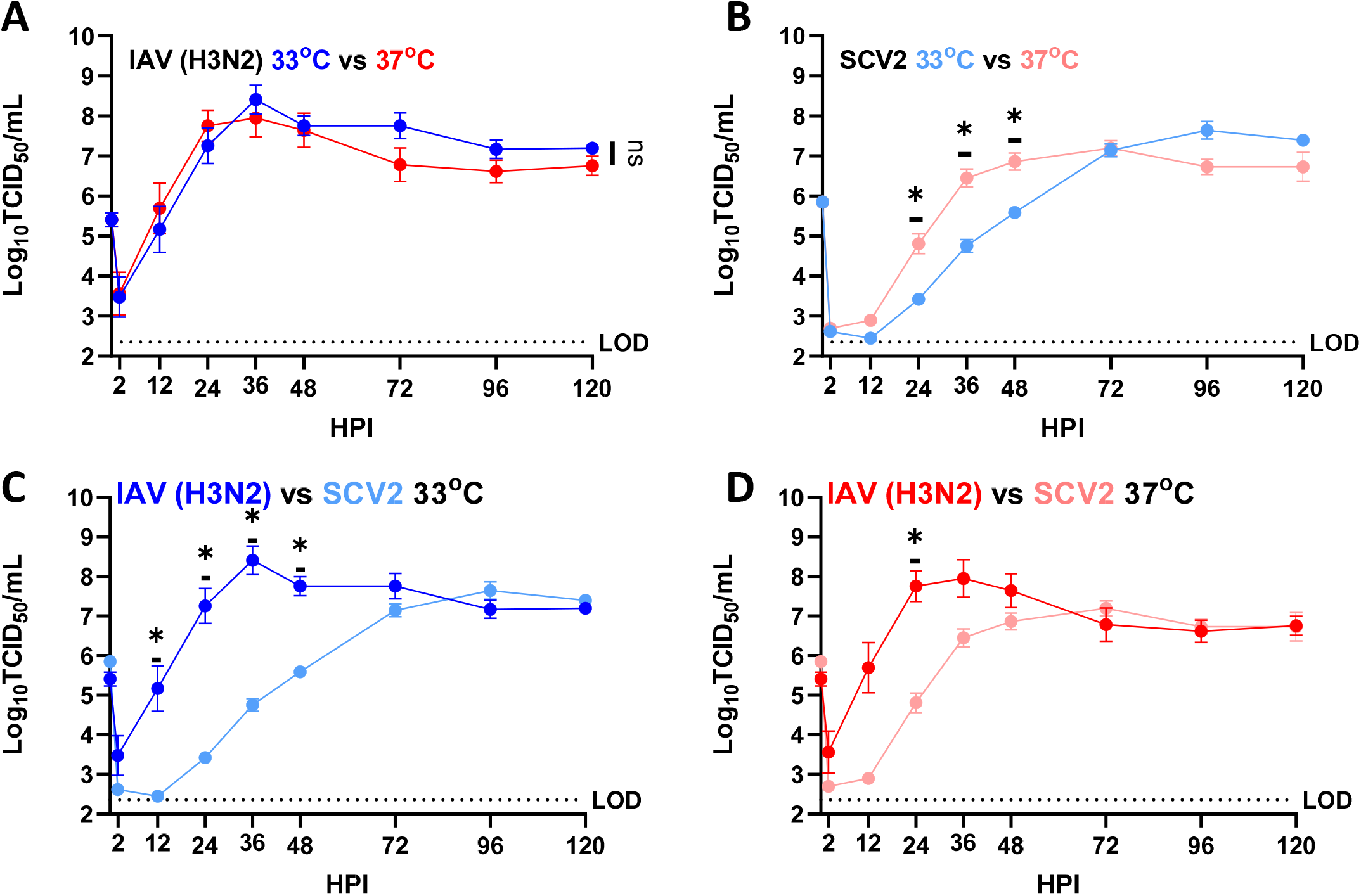
Replication of SARS-CoV-2 and IAV viruses on hNECs at 33°C or 37°C. Multistep growth curves at an MOI of 0.5 infectious units/cell, were performed on hNECs at 33°C and 37°C with the indicated viruses. Data are pooled from two independent experiments each with n=3 wells per virus (total n=6 wells per virus). *p< 0.05 (two way repeated measures ANOVA with Tukey’s posttest, analyzed by timepoint). 0, 2hr excluded from statistical analysis. Dotted line indicates limit of detection. Data is graphed to both show replication of each virus at the two different temperatures (A and B) as well as between the two viruses at the same temperature (C and D).

**Figure 2.**
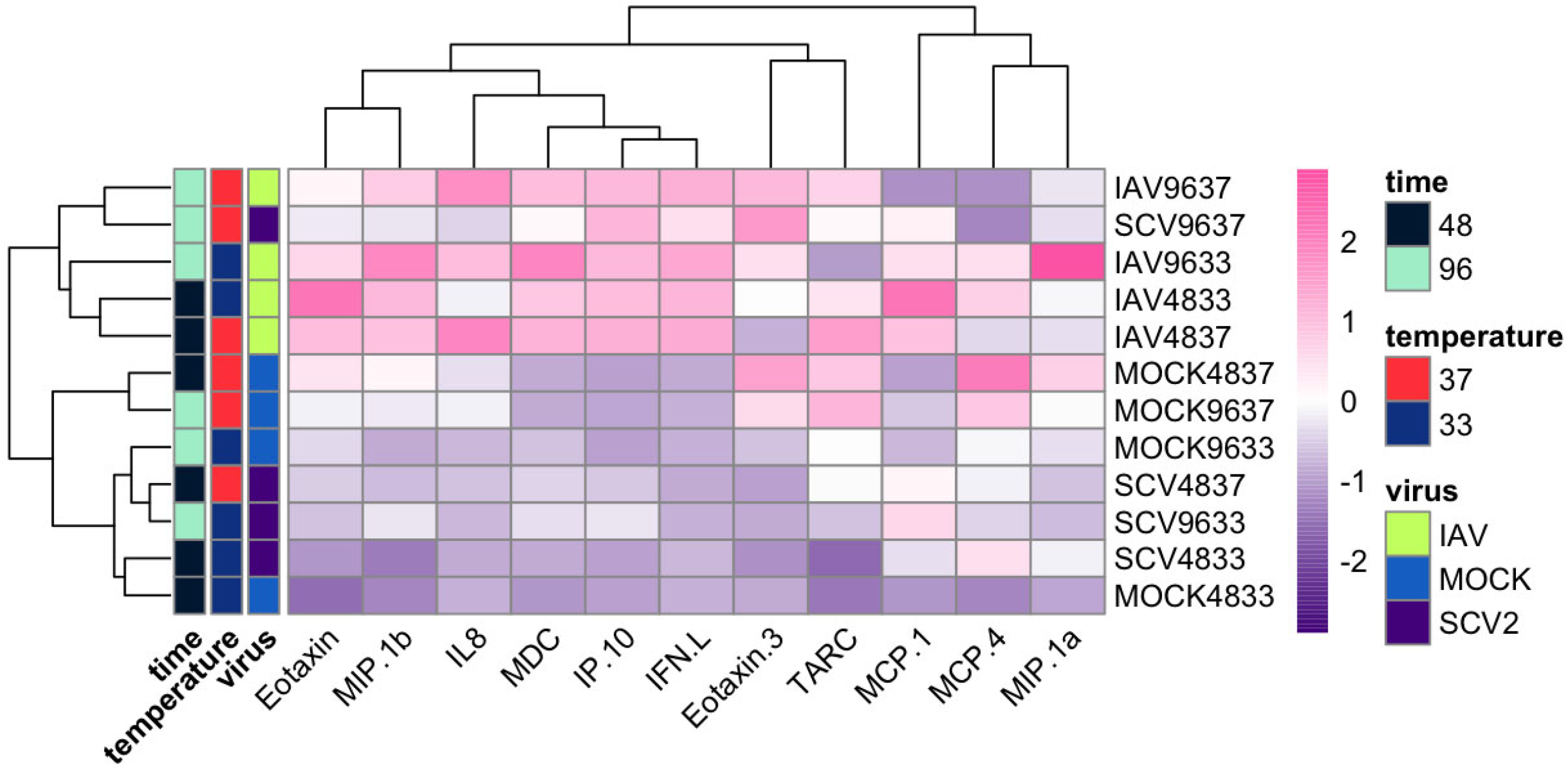
Comparison of cytokine expression induced due to infection at different temperatures over time. Basolateral secretions of cytokines, chemokines, and interferon lambda were measured at 48 and 96 HPI during low MOI multistep growth curve experiment on hNECs at 33°C and 37°C (n=2, 3 wells each, 6 wells total). Values were averaged and then scaled to calculate Z score. Hierarchical clustering was performed based on both analyte and sample.

### RNA-seq of infected hNECs at different physiological temperatures

In order to identify host factors that may be driving both temperature dependent replicative fitness as well as differences in epithelial cell transcriptional responses, RNA-sequencing was performed on mock, IAV, or SCV2 infected hNECs at either 33°C or 37°C. Samples were collected either 24 or 48HPI to focus on early infection responses. The 24HPI samples will be hereafter referred to as “early” while the 48HPI will be referred to as “late” infection samples. Also, 33°C samples will be referred to as “low” temperature while 37°C will be referred to as “high” temperature. Reads were first aligned to the human grch38 genome for annotation, but most late infection samples showed poor (<50%) alignment. Unaligned reads were then aligned to both reference IAV and SCV2 genomes, leading to overall alignment scores of >90% for all samples (Figure 3). Despite low human alignment scores for the highly infected SCV2 and all IAV infected samples, due to the depth of sequencing we were still able to successfully capture responses due to temperature and infection (Supp Figure 1).

**Figure 3.**
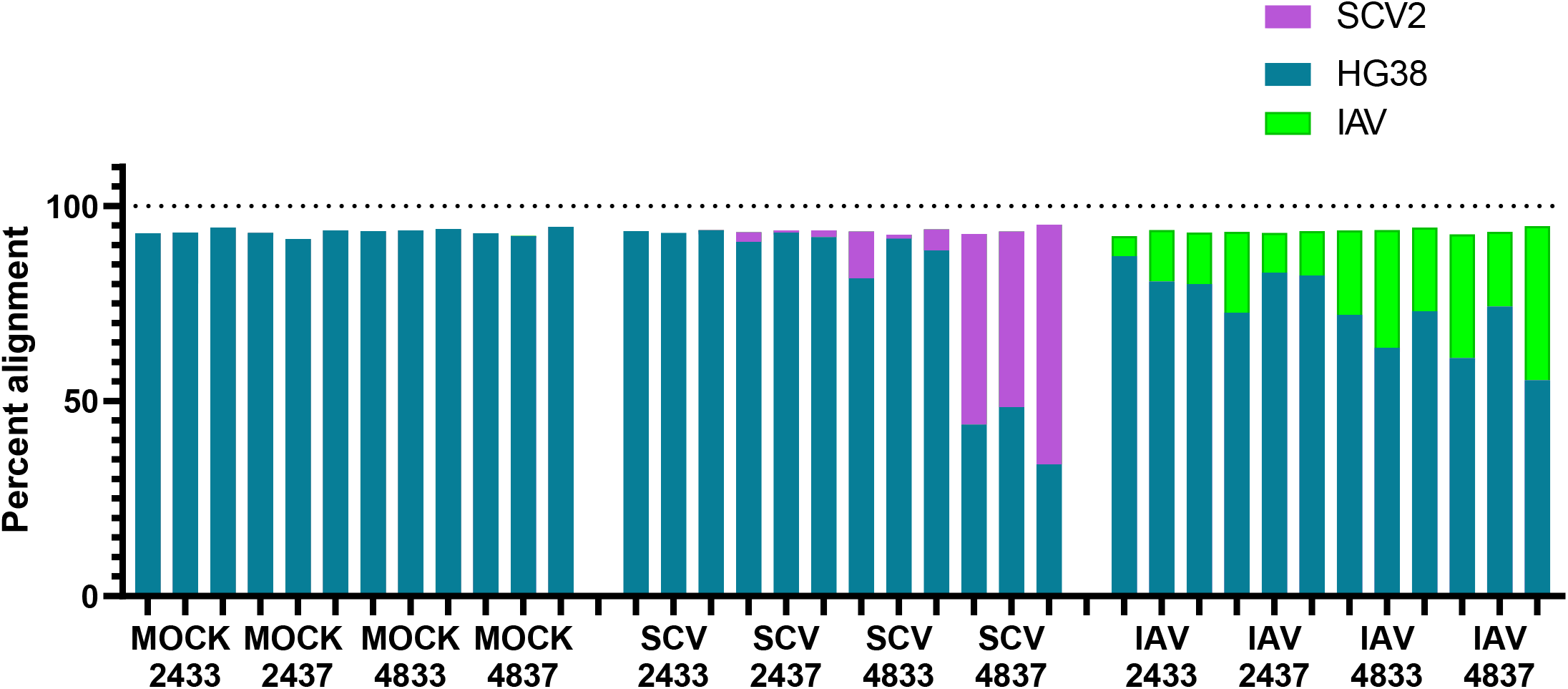
Alignment summary for RNA-seq samples. hNEC cultures were infected with either IAV, SCV2, or mock-infected at 33°C or 37°C and then collected for RNA-sequencing. All sequencing reads were aligned to the human hg38 reference from Ensembl or in-house virus reference sequence downloaded from NCBI. All samples had over 90% alignment to one or more reference sequences. All infected samples had some amount of sequences align to the indicated viral genome suggesting all samples were infected.

### Variance in dataset due to infection and temperature

Differential expression analysis was performed based on temperature, time, and virus, and relative expression data was generated to visualize patterns (Figure 4). IAV infected samples, regardless of time or temperature, seem to cluster separately and most SCV2 infected samples cluster with Mock infected samples. The exception to this is the late, high temperature SCV2 infected samples which cluster with early IAV infected samples. Principal component analysis (PCA) was then performed to identify sources of variance within the dataset (Figure 5A). PC1 explains a majority of the variance in the data (70%). As was seen with relative expression, most late IAV infected samples, regardless of temperature, cluster separately from other samples along PC1. Additionally, most SCV2 samples cluster with mock-infected samples, except for the late, high temperature SCV2 infected which cluster with early IAV infected samples along PC1. The relative expression of the top 12 defining genes was determined to also follow infection state and were mostly related to innate immune responses as well as healthy ciliated cells (Fig 5B). Analysis of the top 500 right PC1 defining and top 150 left PC1 defining genes shows that PC1 describes level of infection response, with the right cluster representing non-responding samples and the left cluster representing high responding samples (Figure 5C and 5D). This is independent of actual infection state, as all samples were determined to be productively infected both by infectious virus quantification in apical wash and by presence of viral genome sequences (Supp Figure 2 and Figure 3).

**Figure 4.**
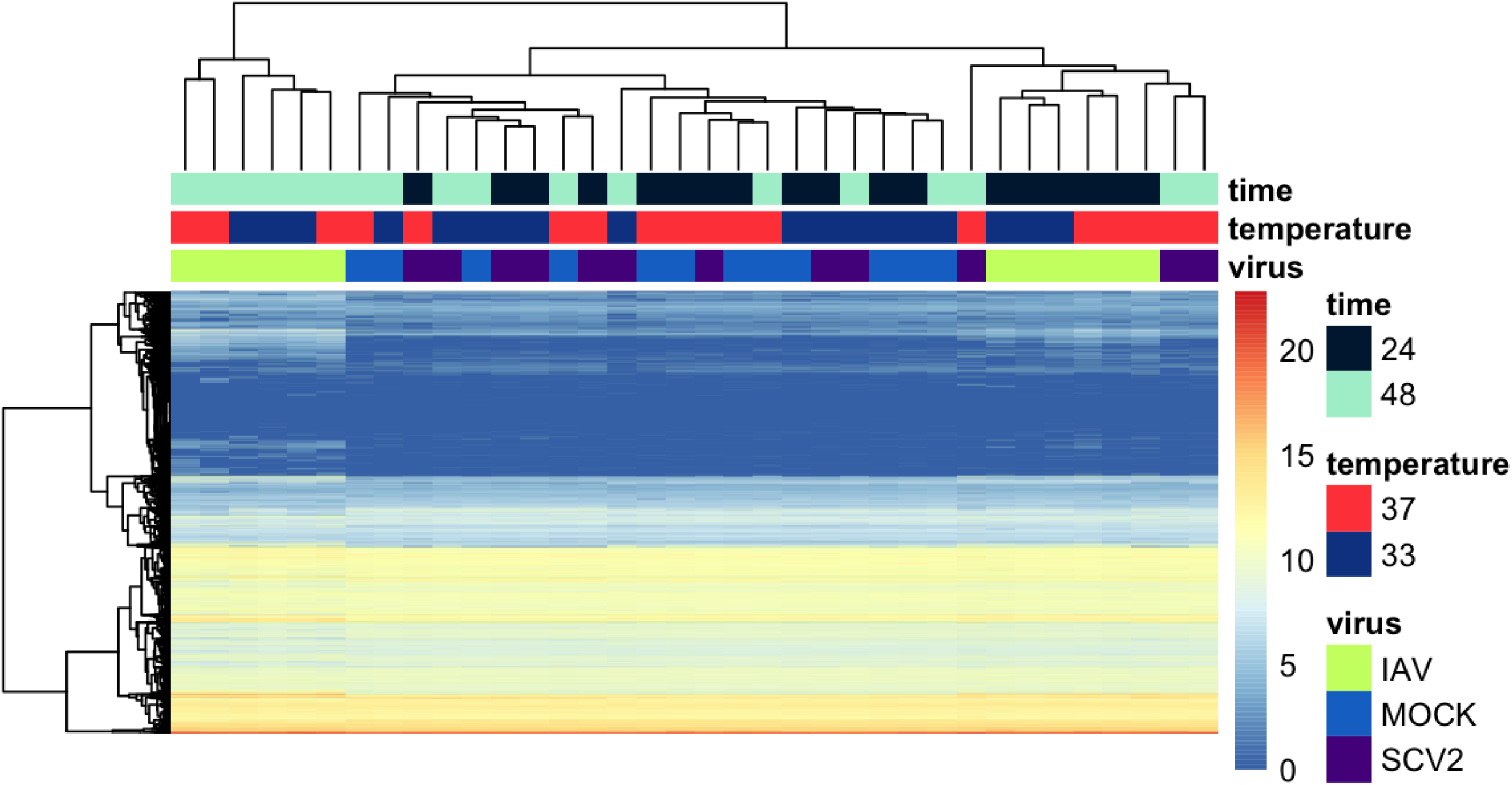
Heatmap summary of gene expression in each condition. Relative gene expression for all genes identified in each sample was calculated and scaled to calculate z score. Hierarchical clustering was performed both by gene and by sample to identify patterns. Samples are labeled by time, temperature, and virus.

**Figure 5.**
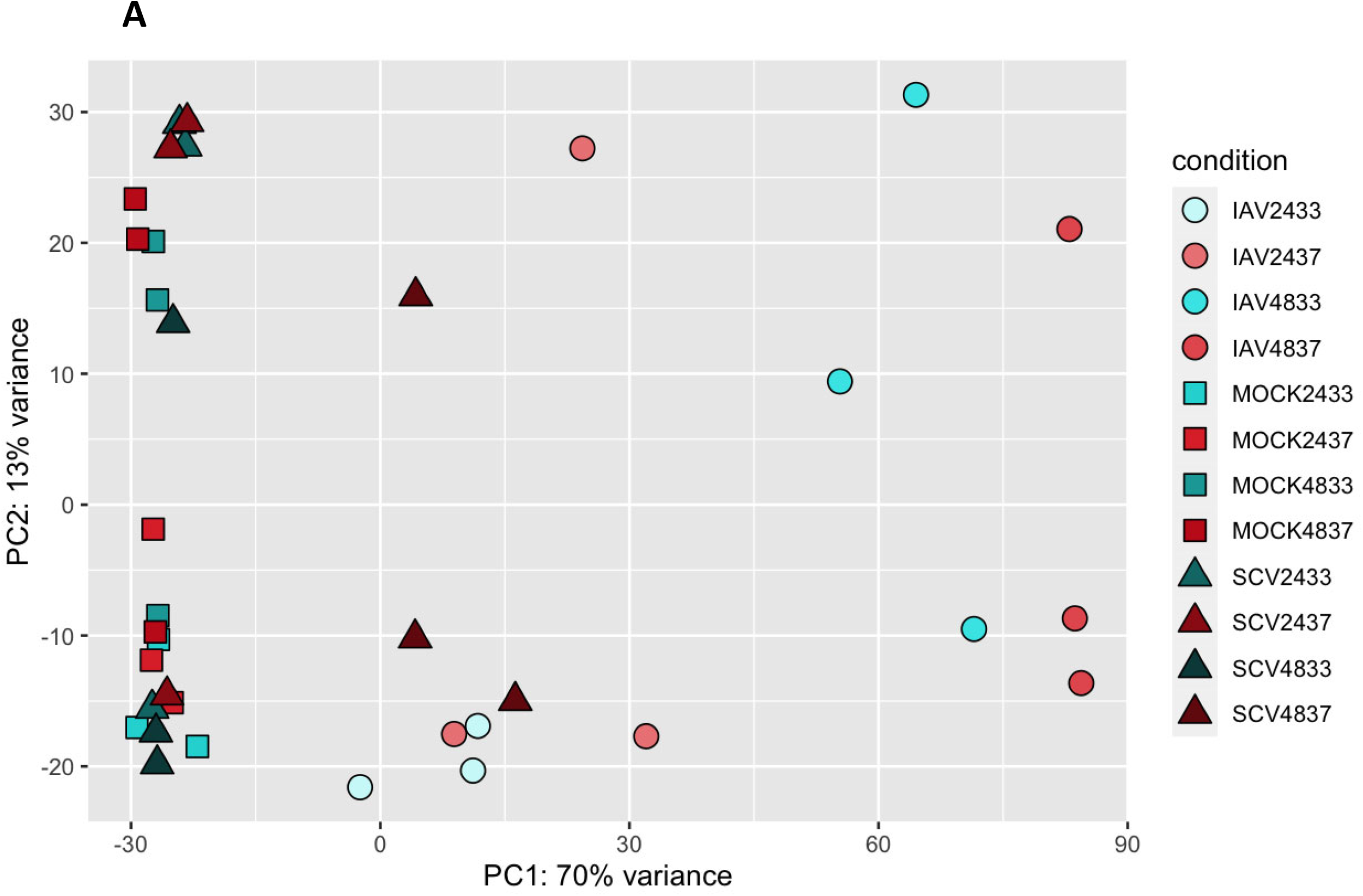
PCA analysis of all RNA-seq samples to identify broad sources of variation. Principal component analysis was run using all expression data for all samples. Samples are colored based on temperature and shape corresponds to condition (IAV, SCV2, or mock-infected)(A). The top 6 genes defining the left and right clusters of PC1 were determined and relative expression in each sample is shown as z score (B). Additionally, the top 500 right defining and 150 left defining genes were used in Biological Process enrichment analysis (C and D). Fraction overlap indicates the number of the top genes that were found to overlap with the indicated pathway divided by the total number of genes in that pathway.

In order to identify variance due to temperature, further principal components (PCs) were investigated. Samples were observed to cluster by temperature along PC4 (Figure 6A). The top 250 genes defining each cluster of PC4 were identified and pathway enrichment revealed that high temperature samples are defined by high infection responses, likely due to the fact that we observe higher levels of infection in these samples (Figure 1, 3, Supp Figure 2, Figure 6B). In contrast, low temperature samples show a strong signature for keratinization (Figure 6C, Supp Figure 5A). The relative expression of the top 17 genes defining the keratinization phenotype was determined and hierarchical clustering confirmed that this is an early marker of low temperature samples regardless of treatment (Figure 6D). Additionally, combined expression of the top 10 differentially expressed keratin-related genes show clear separation between 33oC and 37oC samples (Supp Fig 4).Finally, genes that were more highly expressed in high temperature mock infected samples related to pathways involved in ion transport and tissue development, which may be pointing to metabolic differences between cells at different temperatures (Supp Figure 5B).

**Figure 6.**
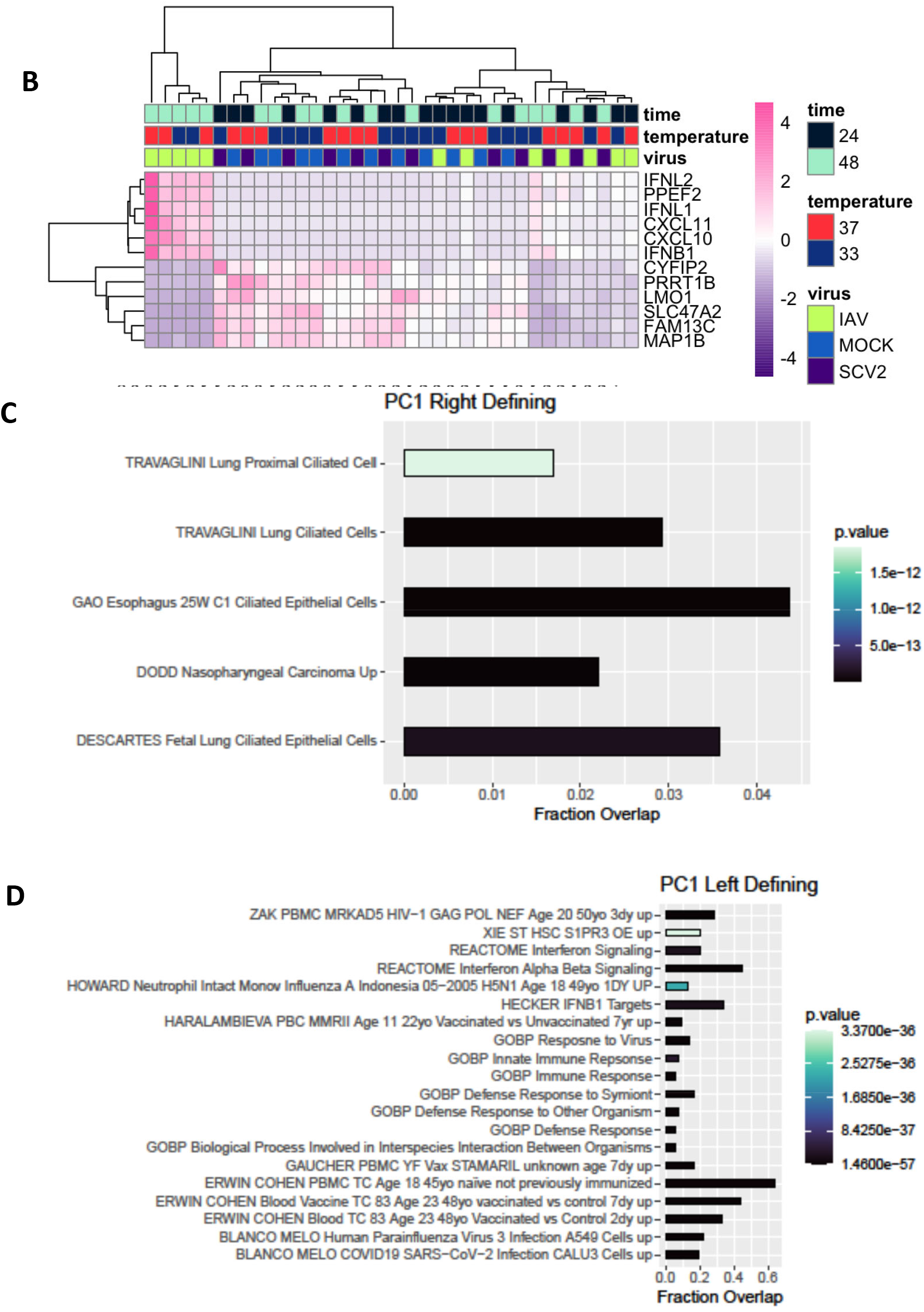
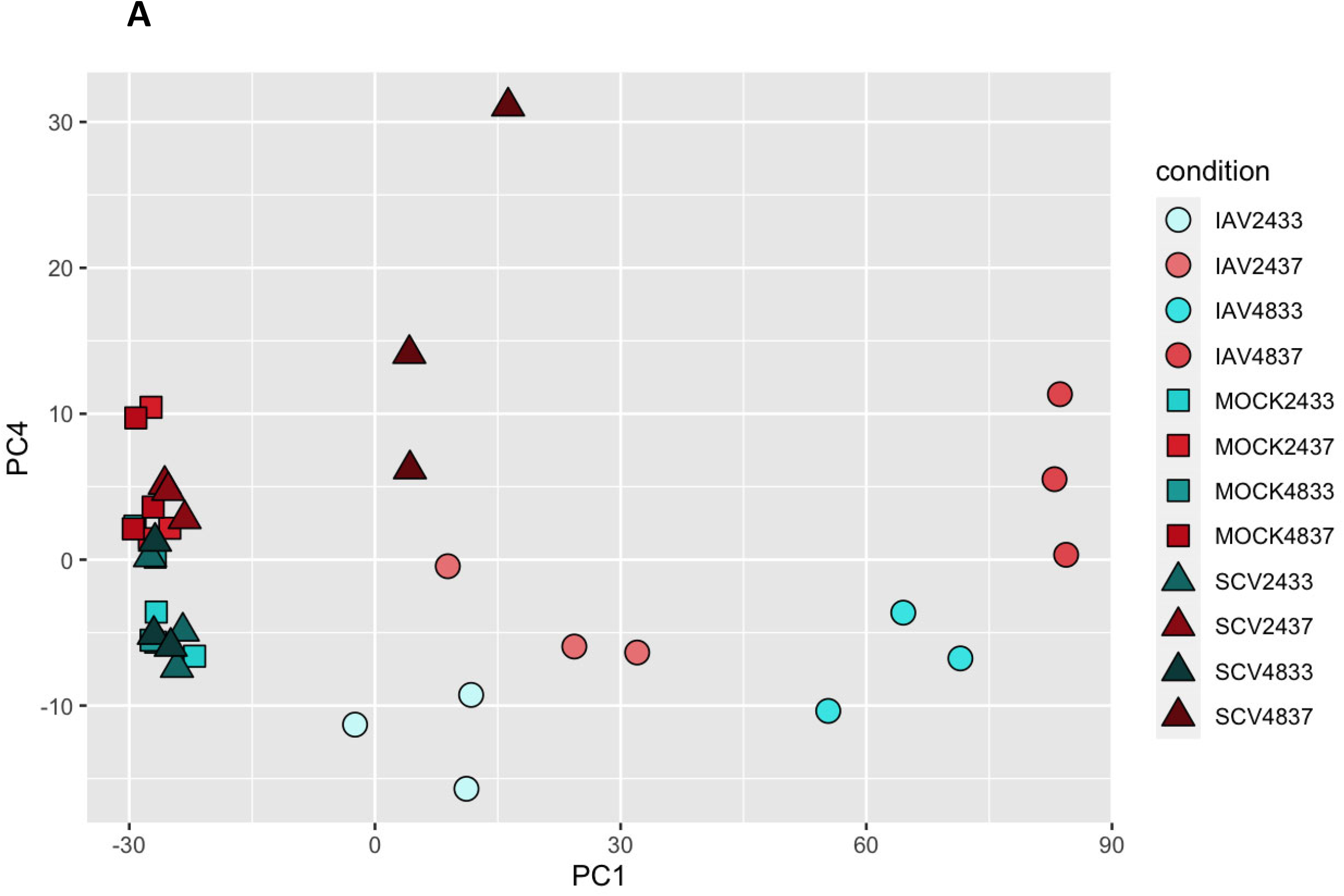
PCA analysis of all RNA-seq samples to identify variation due to temperature. Principal component analysis was run using all expression data for all samples. Samples are colored based on temperature and shape corresponds to condition (IAV, SCV2, or mock-infected)(A). The top 250 right and left defining genes for PC4 were used in Biological Process enrichment analysis (B and C). Fraction overlap indicates the number of the top 250 genes that were found to overlap with the indicated pathway divided by the total number of genes in that pathway. Additionally, the relative expression of the genes defining the keratinization hit in each sample were determined and plotted in a heat map as z score with hierarchal clustering to confirm pattern(D).

### Differentially Expressed Genes (DEGs) and pathway enrichment analysis

We were also interested in differences driven by temperature within each virus infection (IAV, SCV2, or mock) over time. Pairwise comparisons were made for each group at each timepoint and pathway analysis of differentially expressed genes was performed. At baseline (mock-infected), about 100 genes were differentially expressed due to temperature and this number stayed consistent over time (Figure 7A and 7B). Pathway enrichment analysis for these genes mainly pulled out different transcription factors that could be driving this differential expression through epigenetic remodeling, but also identified pathways related to protein binding, organelles, and the cytoplasm (Figure 7C and D). Keratinization also came up as a significant hit when only the top differentially expressed genes were included (Supp Figure 3).

**Figure 7.**
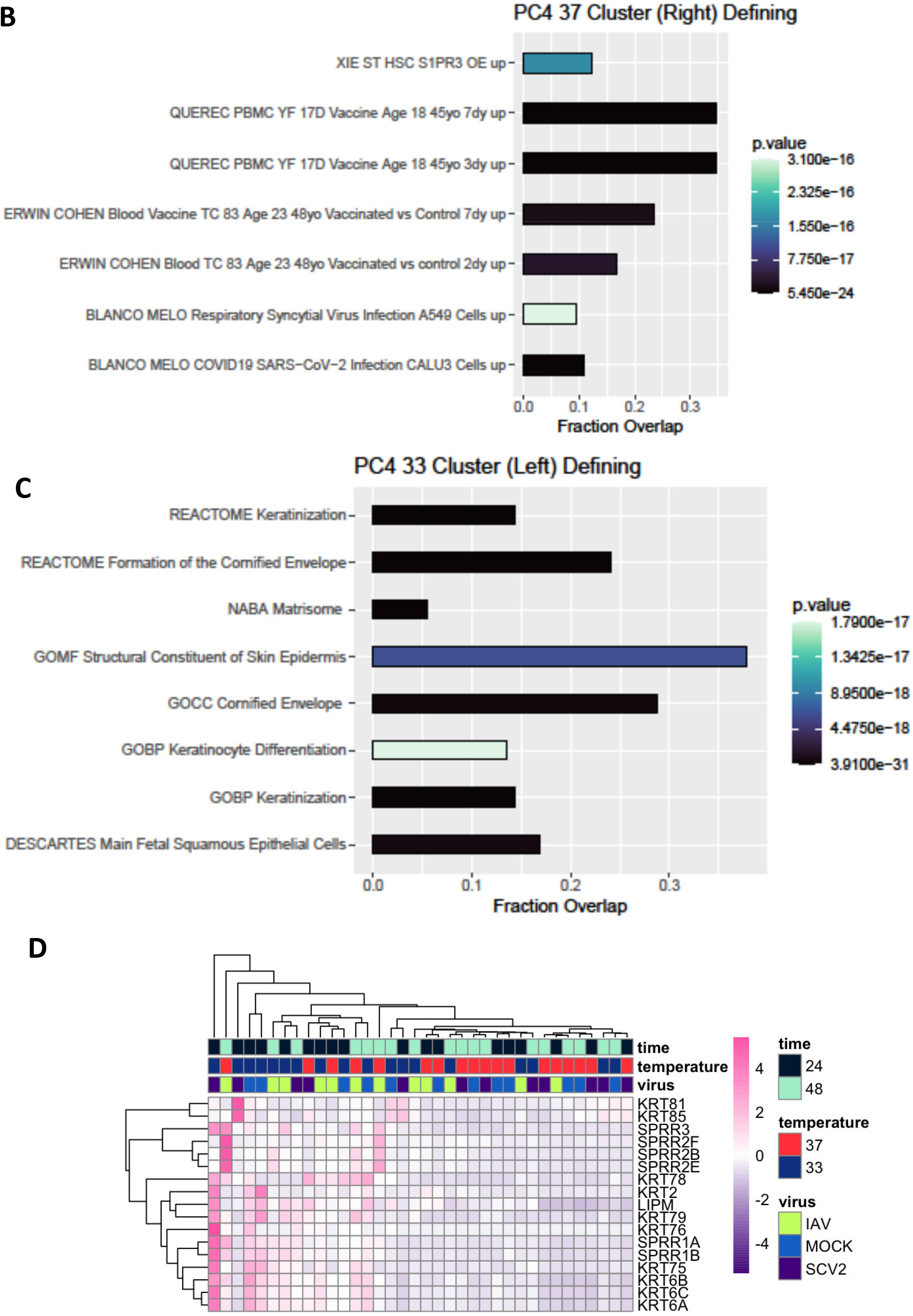
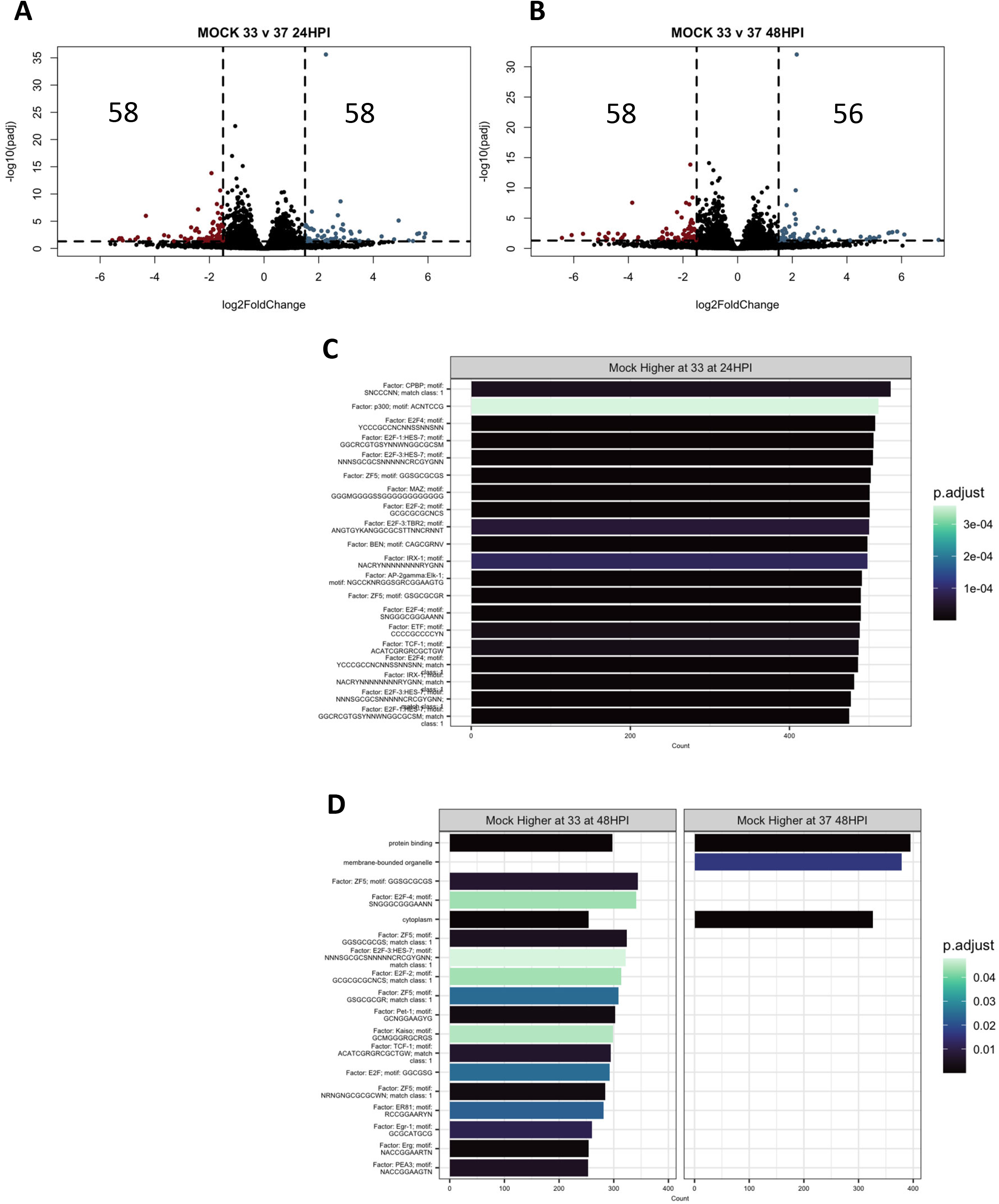
Transcriptomic changes due to temperature over time. Differential expression analysis was run between mock infected samples at 33° or 37°C at 24 (A) or 48 (B) HPI. Significantly differentially expressed genes (p<0.05, log2FC > 0) were then used in pathway enrichment analysis for each timepoint (C, D). The size of the bars indicate the number of genes in that pathway identified, and the bars are colored based on p value.

In order to identify infection-specific temperature differences, each virus infected sample was first compared to its matched mock-infected sample (Figure 8). While IAV infection at any temperature generated a large infection response (Figure 8 A and 8C), only SCV2 infection at high temperature elicited a significant transcriptomic response (Figure 8B and 8D). Due to this, the low temperature SCV2 samples were not used as an independent condition for analysis. IAV infection at different temperatures showed moderately different transcriptomic profiles, with an increase in the number of differentially expressed genes over time (Figure 9A and 9B). At the early timepoint, most differentially expressed genes at 37°C were in pathways related to signaling in response to various stimuli (Figure 9C). At the late time point, most differentially expressed genes at 33°C were in pathways related to biological regulation and homeostasis as well as signaling through vesicles and junctions (Figure 9D). This suggests that while the baseline antiviral response at different temperatures remains consistent, there may be metabolic or signaling changes that affect how well the cultures are able to adapt to new pressures such as infection.

**Figure 8.**
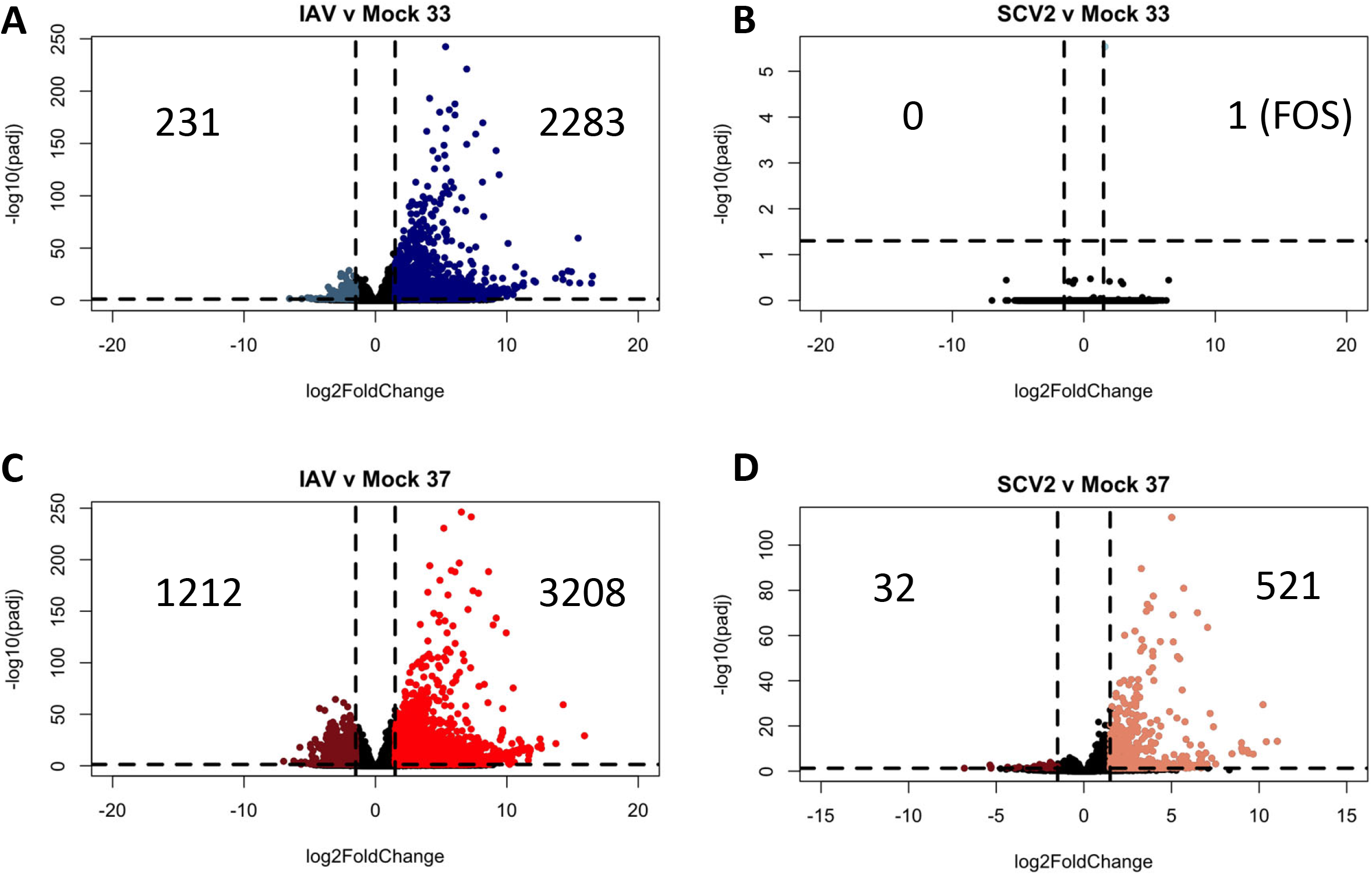
Differential gene expression due to infection. Differential expression analysis was run between infected samples at 33° or 37°C at 48HPI and their matched mock infected sample as indicated (A-D). Data were pooled from three replicate wells. Log 2 fold change indicates the mean expression for a gene. Each dot represents one gene. Black dots indicate no significantly differential expression between the two indicated groups. Colored dots indicate both a significant p value (padj <0.05) and log 2 fold change (1.5).

**Figure 9.**
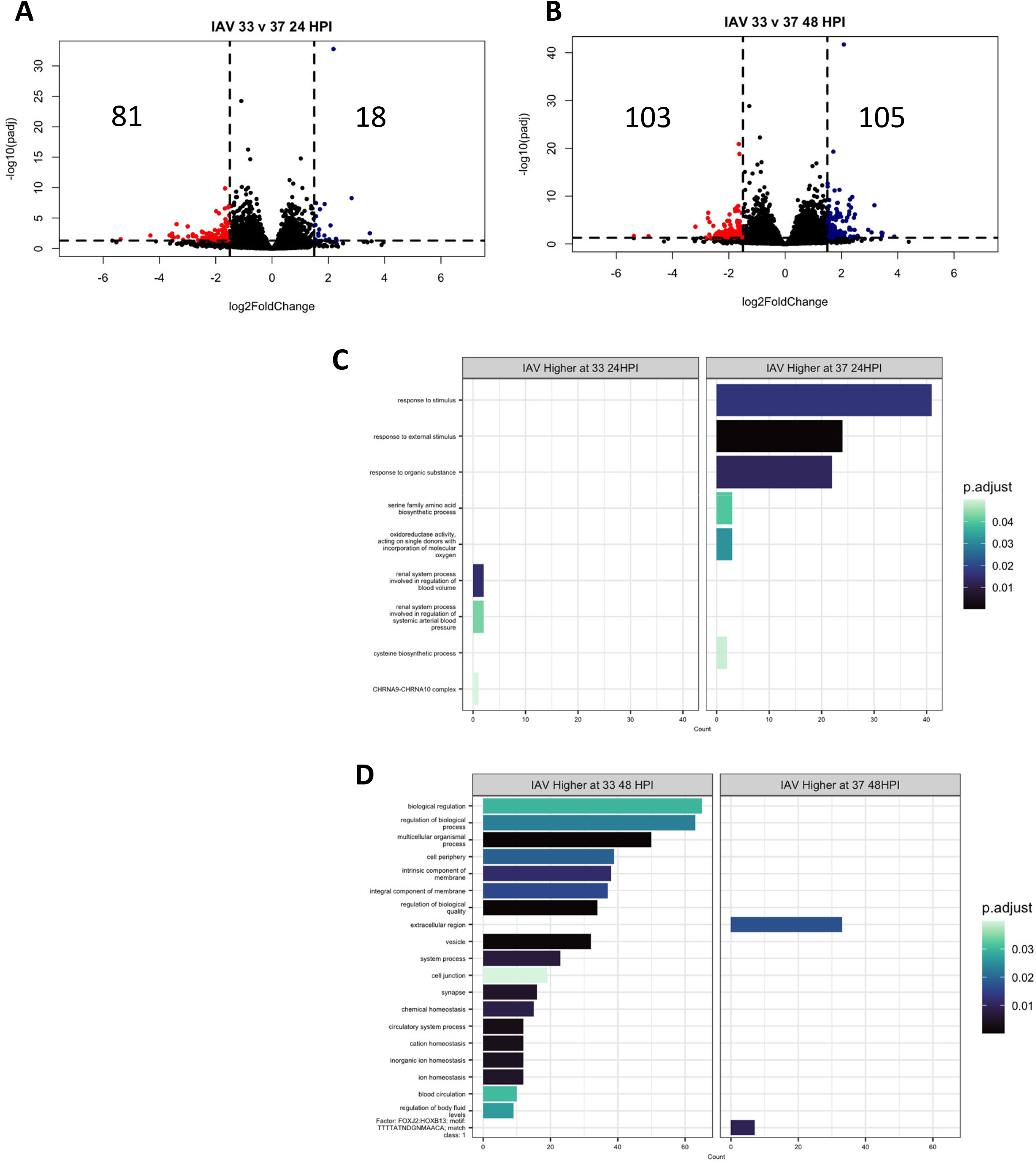
Transcriptomic changes due to temperature over time. Differential expression analysis was run between IAV infected samples at 33° or 37°C at 24 (A) or 48 (B) HPI. Significantly differentially expressed genes (p<0.05, log2FC > 1.5) were then used in pathway enrichment analysis for each timepoint (C, D). The top 30 hits are shown. The size of the bars indicate the number of genes in that pathway identified, and the bars are colored based on p value.

To identify whether there are differences in the epithelial cell response due to virus used, high temperature, late timepoint SCV2 and IAV infected samples were compared (Figure 10). Overall, there were almost 3000 differentially expressed genes due to virus infection (Figure 10 A). IAV infected samples showed increased expression of genes in pathways related to the membrane and membrane associated factors, while SCV2 infected samples showed increased expression of genes related to cell projections (Figure 10 B and 10C). This difference could be due to slightly different replication cycles used by IAV and SCV2 viruses or could reflect how the cultures attempt to heal from infection, as the 48HPI timepoint is peak production of IAV infectious virus.

**Figure 10.**
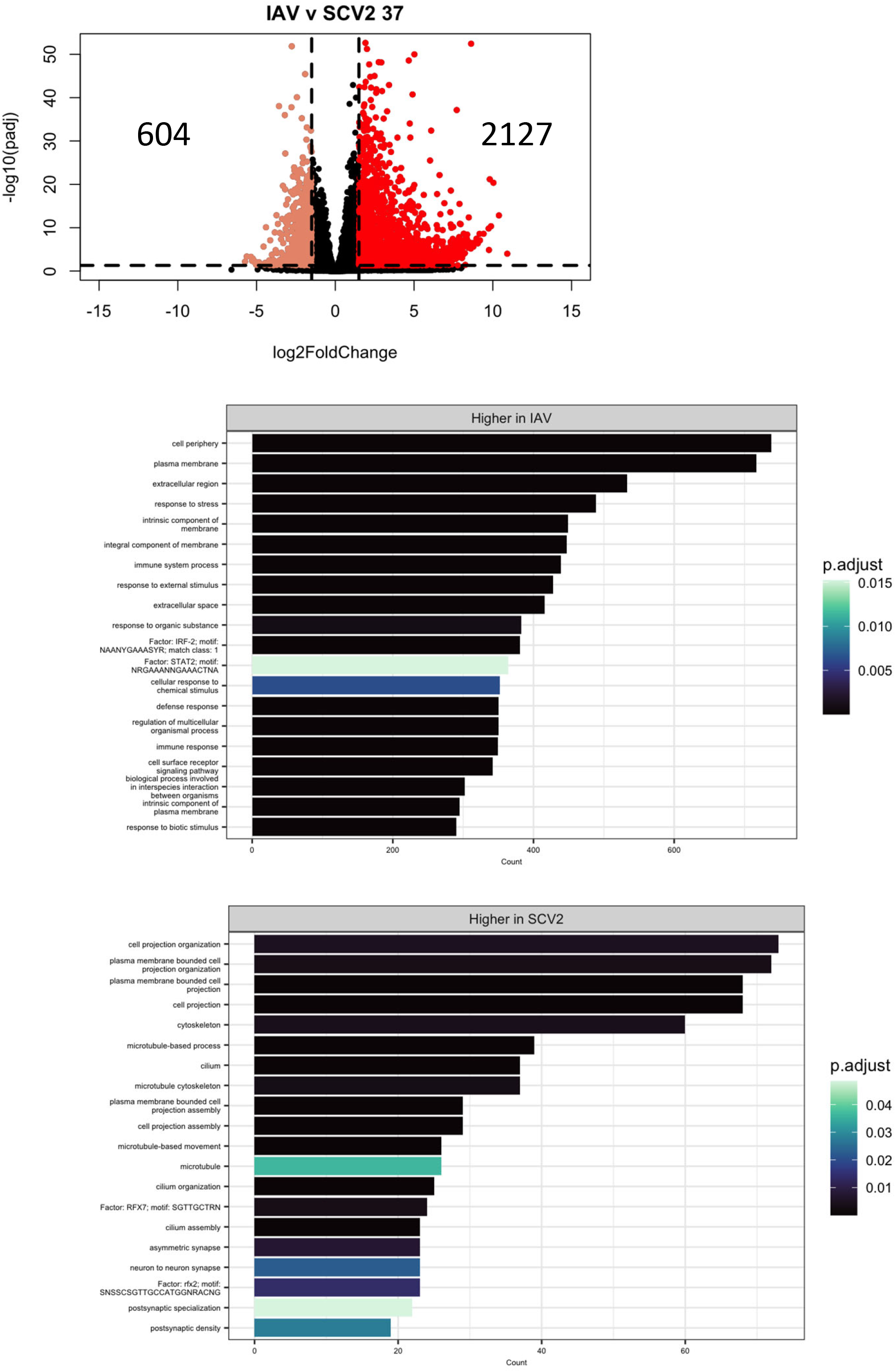
Transcriptomic changes due to virus used for infection. Differential expression analysis was run between IAV and SCV2 infected samples at 37°C 48HPI (A). Significantly differentially expressed genes (p<0.05, log2FC > 1.5) were then used in pathway enrichment analysis for each timepoint (B, C). The size of the bars indicate the number of genes in that pathway identified, and the bars are colored based on p value.

Finally, to identify genes that drive temperature specific infection responses, differentially expressed gene lists due to temperature within each condition (IAV, SCV2, mock infection) were compared (Figure 11). The lists were generated from the pairwise comparisons (Figure 7B, 9B, SCV2 not shown). Overall, SCV2 infection at 37°C was the most unique with 611 DEGs, likely due to the fact that SCV2 infected samples at 33°C are highly similar to mock. Both SCV2 and IAV infection at 37°C uniquely had higher expression of 18 genes in common, including APOBEC3 genes (APOBEC3A, APOBEC3B, and APOBEC3B-AS1), ssDNA binding protein SHOC1, and pseudogenes CLCA3P and NCF1B, suggesting higher chromatin instability during infection at higher temperatures (Supp table 1). High temperature infections also uniquely upregulated immune genes such as CCL20 and AIM2. Additionally, high temperature infections uniquely upregulated STX19, which is involved in SNARE binding and could impact the efficiency of viral fusion ^20^. In contrast, immune related genes such as IFITIM10 and MPV17L were some of the 14 genes uniquely upregulated due to IAV or SCV2 infection at lower temperatures (Supp table 2). All high temperature samples (IAV, SCV2, and mock) uniquely had higher expression of three genes-CLCA3P, RNAS1, and SLC51B-two of which are involved in transport (Supp table 3). In contrast, low temperature samples all uniquely had higher expression of four genes-CYP4A11, TGFBR3L, RNF32-DT, and RBM3-three of which are involved in metabolism and proliferation (Supp table 4).

**Figure 11.**
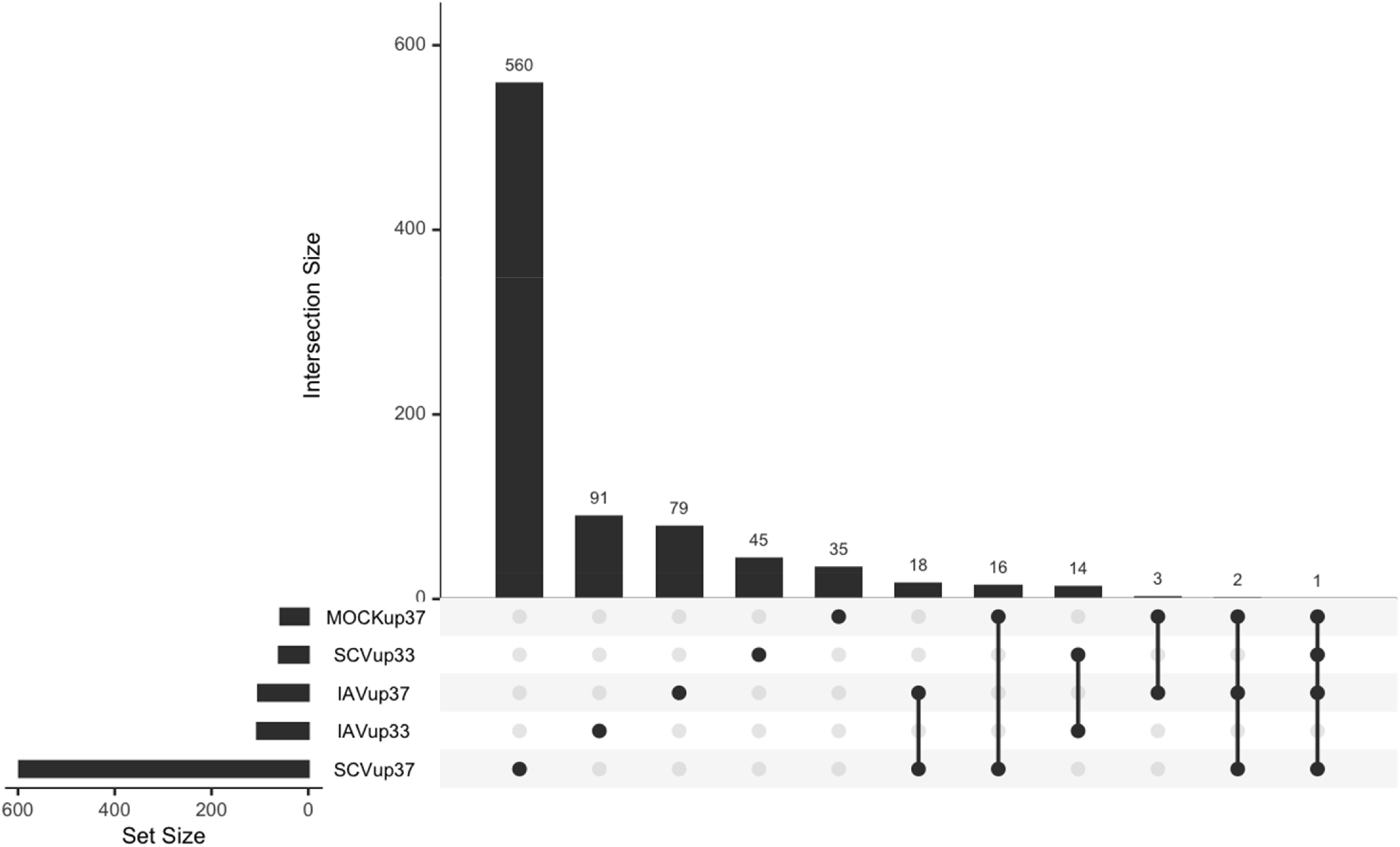
Comparison of genes differentially expressed due to temperature in each condition. Differential expression analysis was run with each condition between 33°C and 37°C 48HPI. Significantly differentially expressed genes were defined as p<0.05 and log 2 fold change of 1.5. Lists of differentially expressed genes were generated and used to create an upset plot. Set size is the number of genes in the indicated category. Intersection size is the number of overlaps for the comparison indicated. Lists corresponding to each intersection are available in supplement.

## DISCUSSION

Both Influenza A and SARS-CoV-2 viruses are respiratory pathogens responsible for some of the most severe pandemics in modern history and remain a threat to public health today^1,2,3,9^. Understanding how these viruses interact with the nasal epithelium -the initial site of infection-while accounting for the effects of physiological temperature is imperative to not only developing intervention strategies, but also understanding how initial infection can transition into severe disease, characterized by progression of infection from the upper to the lower respiratory tract ^11^. While IAV and SCV2 share many similarities, the two use different receptors for entry leading to different cell tropism within the respiratory epithelium ^4,12,18,19,22,23,24,25^. This may influence cellular responses to infection and impact temperature sensitivity due to the location of susceptible cell types along the respiratory tract^20^.

Many studies to date have investigated differences in transcriptional responses to IAV and SCV2, a subset of which are cited here^26,67,81,82^. However, a majority of studies have focused on easily accessible patient blood samples for biomarker discovery, or take samples from deceased patients which represent late infection timepoints and heterologous cellular samples that are more difficult to draw specific mechanistic conclusions from ^14,15,81,82,83^. Additionally, responses to infection have been shown to be cell specific, therefore great care must be taken to be specific about the context of conclusions drawn from these types of studies^20^. Even within respiratory epithelial cells, it is important to distinguish between cultures derived from the upper and lower respiratory tract when designing these studies, as there are important differences in cell type proportion and receptor expression to account for in addition to specific microenviron-ments^61,62,63^ Other groups have reported an attenuated response to early SCV2 infection, especially in nasal epithelial cells, with most responses not being observed until 72HPI^26,67^. This study has important limitations, namely the use of a single donor for the RNA-sequencing samples to limit variability so the focus could be on specific virus and temperature-related responses. Also, the use of a bulk RNA-seq approach may mask some cell-type specific responses. However, this approach will reveal average epithelial changes that are more likely to be targetable in general disease conditions^84,85^.

In order to understand if physiological temperature influences IAV and SCV2 replication, growth curves were performed at the extremes of the range of normal respiratory tract temperature ^27,28,29^. SCV2 showed more sensitivity to lower temperatures, leading to slower replication kinetics, while IAV was not affected. Additionally, infection with IAV induced a larger innate immune response earlier than infection with SCV2, and overall innate immune induction was observed to be higher at higher temperatures. It is unclear whether the delayed innate immune response in SCV2 infected cultures at 33C represents a delayed induction or enhanced active suppression of innate immune response. Comparison of 24 HPI IAV infected samples to 48HPI SCV2 infected samples show less differentially expressed genes than matched timepoint samples but still contain significant differences (supp fig 6). Delayed or reduced interferon responses have previously been identified as risk factors for severe disease, which may explain the higher morbidity rates observed in SCV2 infection cases compared to IAV infection ^10,14^.

Transcriptomic changes in nasal epithelial cell cultures revealed that both temperature and virus influence the host infection response. One of the most striking phenotypes was a strong keratinization signature observed at 33°C regardless of infection state. Keratin proteins are most often produced from suprabasal cells in the epithelium and can assist in the integrity and mechanical stability of epithelial cell to cell contacts as well as in single cells ^35,36^. They also play important roles in signaling, transport, and growth ^37^. Two of the keratins identified-76 and 78-are found predominately in the palate and tongue, suggesting that the difference in temperature may impact differentiation patterns and help form the structure of the respiratory tract ^37^. Alternatively, keratinization also plays a role in wound healing, and may be a sign that the cells are able to recover from viral infection damage quicker, which may explain why there is greater persistence of infection at lower temperature with the maintenance of newly differentiated, susceptible cell types ^38^. It has also been associated with the damage response in the nasal epithelium due to cigarette use^87^. Hyper-keratinization can also be a response to chronic irritation, again a likely response to increased persistence of infection at lower temperatures ^38^. A study investigating asthma-mediated protection from SCV2 infection reports higher keratinization of the cells, mediated through IL-13 signaling, to be a mechanism of protection against high viral loads^86^.However, more research is needed to identify how temperature impacts keratinization, as well as the impact of keratinization on viral replication.

It is interesting to note that both lower temperature and infection with SARS-CoV-2 virus appears to delay responses to infection. RNA-seq results show limited cellular responses to infection with SARS-CoV-2, especially when infected at 33°C. It is worth wondering whether the response to SARS-CoV-2 infection is the same as the response to IAV infection but delayed. Measuring the innate response via cytokine and chemokine production 48 and 96 HPI suggests that the SCV2 37°C response eventually become detectable, suggesting that the delay persists until responses are more likely being driven by damage-associated molecular patterns (DAMP) rather than pathogen-associated molecular patterns (PAMP) signaling^71^. SCV2 inhibition of IFN signaling has been reported extensively in the literature and has been hypothesized to be a mechanism for increased transmissibility of variants^10,14,32,33,34,70^.Additionally, the clusters generated using the cytokine data seem to be driven by both temperature and time. This is especially interesting in the IAV condition where there is no difference in the amount of infectious virus produced over time, but there are differences in cellular response due to temperature that are most obvious at 96HPI. This is in contrast to other literature that shows innate responses being proportional to infectious virus titer and warrants further study^69.70^.

Although IAV and SCV2 are both respiratory pathogens, SCV2 has broader cell tropism and the two viruses have different mechanisms for replication^4,12,18,19,22,23,24,25^. These differences can impact cellular responses to infection^20^. We found that the response of the hNEC cultures to IAV infection was driven by genes involved in pathways related to membrane and cell signaling, while the SCV2 response was driven by genes related to the cytoskeleton and cellular projections. Recent work has shown that SARS-CoV-2 manipulates the cilia and microvilli of epithelial cells in order to initiate infection and spread to neighboring cells ^39^. Our data suggests that this is unique to SARS-CoV-2 infection, and that IAV in contrast uses membrane transport systems for entry and transport of viral factors to the plasma membrane where viron assembly occurs^40^. The differences in pathways used for viral movement within a cell could give insight into cell type targets of each virus separate from receptor expression, identifying other host factors needed for productive infection^66^. Additionally, identifying the differences in how these viruses manipulate the host cell to allow for replication and release could also help to explain the differences seen in pathogenesis between these viruses, and identify new targets for intervention.

In addition to differences being seen between distinct respiratory pathogens, different SCV2 variants have been observed to have different sensitivity to temperature ^72^. This study was performed with an ancestral SCV2 variant, which along with Delta variant viruses (defined by the spike protein) has been shown to primarily target the lower respiratory tract^72,73^. These variants have also been shown to be more sensitive to lower temperatures^72,73^. In contrast, Omicron variant viruses do not have the same sensitivity to temperature that has been observed in prior variants and have been observed to target the upper airway^72,79^. This may be due to differences in entry pathway preferences and cellular tropism^75,76,77^. The location within the respiratory tract where the virus replicates has important implications for clinical outcomes^80^. Future work will investigate viral factors that contribute to temperature sensitivity. For example, in IAV infection, mutations in both the HA and M2 protein, along with the replication machinery, have been shown to impact temperature sensitivity in subtype specific manners^68,69^. In SCV2, the non-structural proteins have been implicated in manipulating the host response to infection, making them enticing targets to look at in regards to temperature sensitivity^78^. It is likely there is a combination of entry and internal factors that impact viral temperature sensitivity, but this remains an open area of study.

Finally, we identify genes that are commonly differentially regulated due to temperature and infection with IAV or SCV2 viruses. These genes are related to genome defense and general immune responses, as well as membrane fusion, and represent a common cellular target that could be used for treatment of both IAV and SCV2 infections. The vast majority of interferon and innate immune pathways were not expressed in a temperature dependent manner, suggesting a temperature independent induction of these antiviral pathways. Other studies have also shown that SCV2 replication is inhibited at elevated temperature as well, such as those reached during the fever response, again independent of the IFN response^74^. Further research should be done to see if these genes are common to all respiratory pathogens, or that it just happens to be similar for the viruses tested. Additional research is needed to identify each host protein’s role in temperature-dependent viral replication.

Taken together, these data indicate that temperature should be accounted for when evaluating human pathogens and can assist in identifying new treatment strategies as well as understanding the basic biology underlying respiratory virus infection of epithelial cells.

## Author Contributions

Conceptualization, A.P. and J.D.R.; methodology, A.P. and J.D.R.; formal analysis, J.D.R. and M.A.B.; resources, A.P. and M.A.B; data curation, J.D.R.; writing–original draft preparation, J.D.R., M.A.B., and A.P.; writing–review and editing, J.D.R., M.A.B., and A.P.; visualization, J.D.R., and M.A.B.; supervision, A.P.; funding acquisition, A.P. and M.A.B. All authors contributed to the article and approved the submitted version.

## Data Availability Statement

All RNAseq data is available at NCBI Sequence Read Archive, NCBI BioProject: PRJNA925547 and analysis code is available on https://github.com/JRes9/Resnicketal_IAVvSCV2temperature_2023.

## Acknowledgements

We thank the members of the Andrew Pekosz, Sabra Klein, and Kimberly Davis laboratories for insightful comments and discussions pertaining to this manuscript. This work was supported by T32GM007814-37 (JR), T32AI007417 (JR), NIH R01 HG012367 (MAB), the Centers of Excellence for Influenza Research and Response (NIAID N272201400007C) and the Richard Eliasberg Family Foundation (AP). We thank the Bloomberg Flow Cytometry and Immunology Core for use of MSD instrument. We also thank Anne Jedlicka and Amanda Dziedzic of the Johns Hopkins Bloomberg School of Public Health Genomic Analysis and Sequencing Core Facility for their help with preparing and sequencing samples.

