## Supplementary material for "Early transcriptional responses of human nasal epithelial cells to infection with Influenza A and SARS-CoV-2 virus differ and are influenced by physiological temperature": supp figure 2

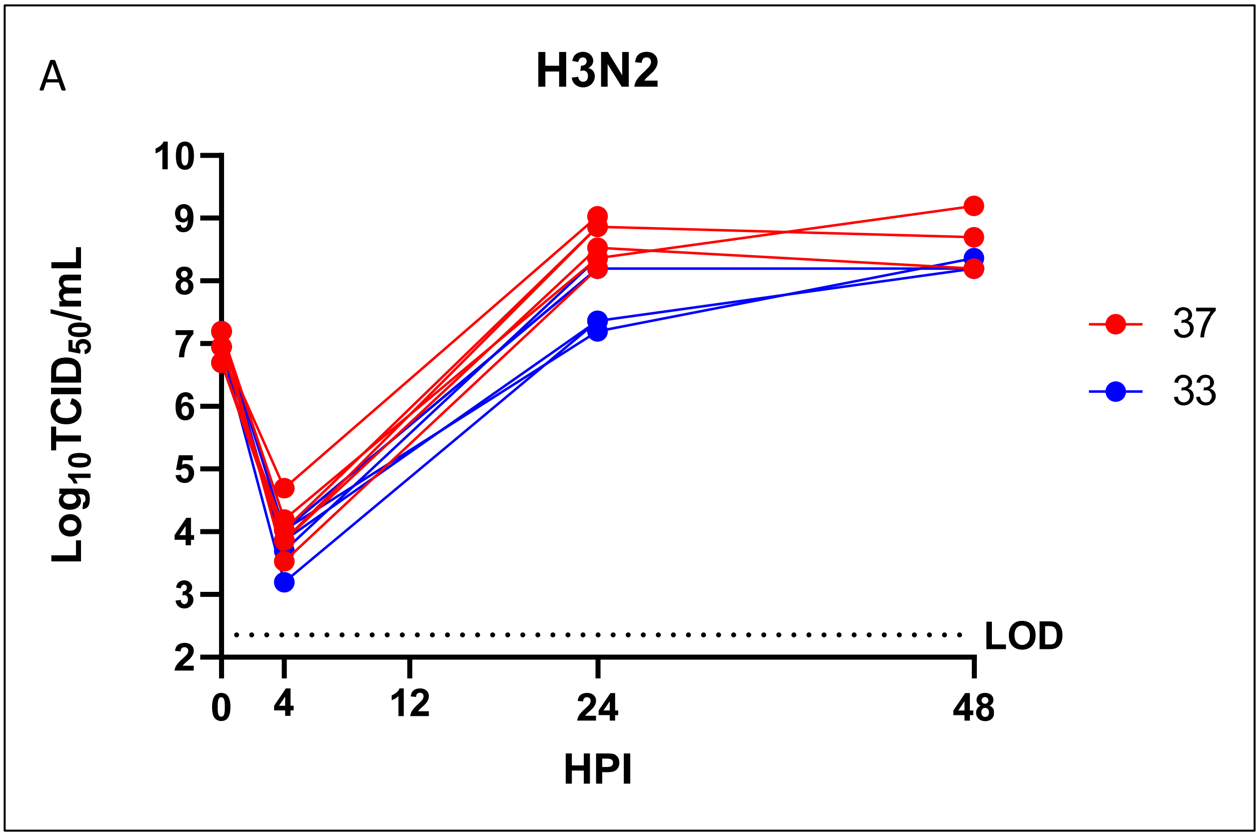


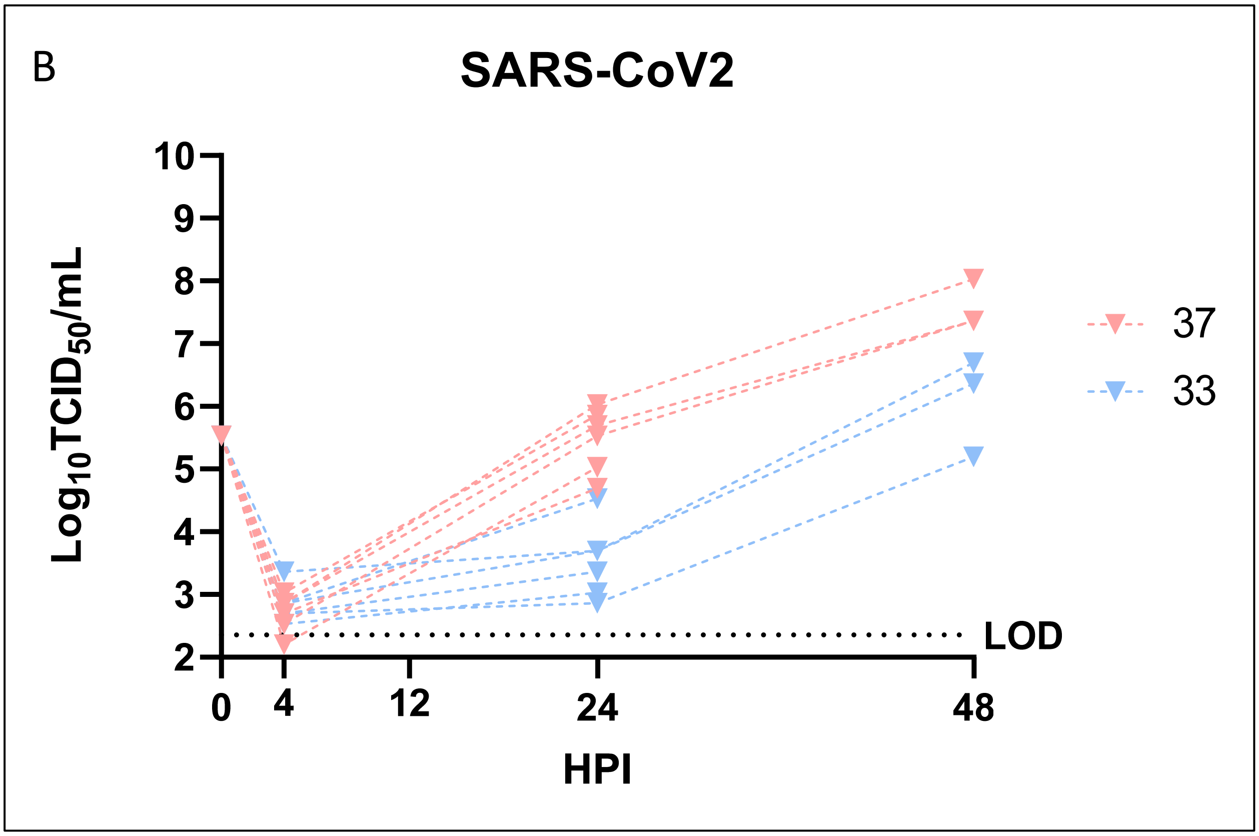


**Supp Fig 2:** *Infectious virus present in apical washes of RNA-seq samples infected with SARS-CoV-2 (B) and IAV (A) viruses at 33oC or 37oC.* hNEC cultures were infected at low MOI, and followed for a multistep growth curves at 33ºC and 37ºC with the indicated viruses. Apical washes were taken at the timepoint indicated before sample collection. Each sample is graphed individually, and 48hr samples were also checked at 24HPI.
