## Supplementary material for "Early transcriptional responses of human nasal epithelial cells to infection with Influenza A and SARS-CoV-2 virus differ and are influenced by physiological temperature": supp figure 3

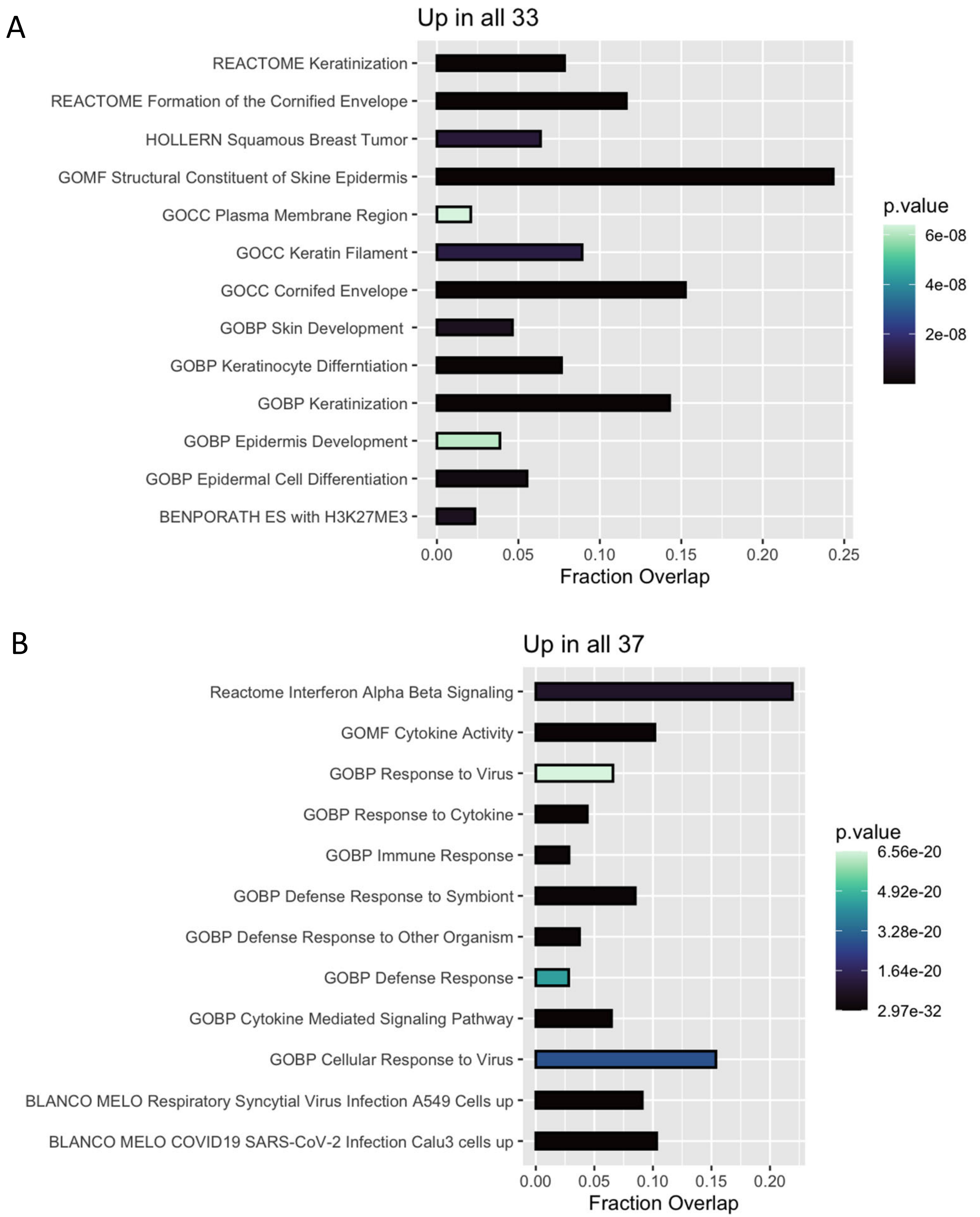


**Supp. Fig 3.** *Pathway analysis of differentially expressed genes between all 33°C and 37°C samples.* Differential expression analysis was run between all 33 and 37°C samples, regardless of virus treatment, and the top 250 genes were used for pathway enrichment analysis.
