## Supplementary material for "Early transcriptional responses of human nasal epithelial cells to infection with Influenza A and SARS-CoV-2 virus differ and are influenced by physiological temperature": supp figure 4

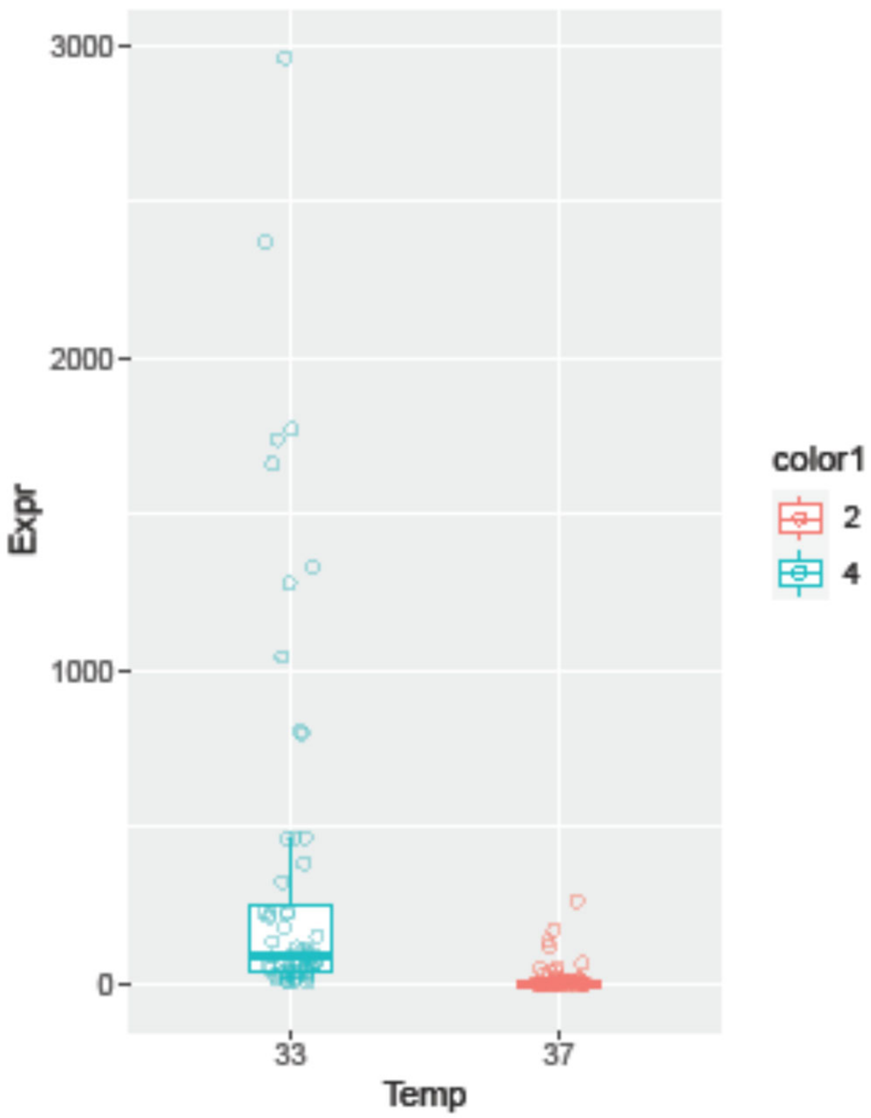


Supp Fig 4: Combined expression of the top 10 most differentially expressed genes in all samples due to temperature.
