## Supplementary material for "Early transcriptional responses of human nasal epithelial cells to infection with Influenza A and SARS-CoV-2 virus differ and are influenced by physiological temperature": supp figure 5

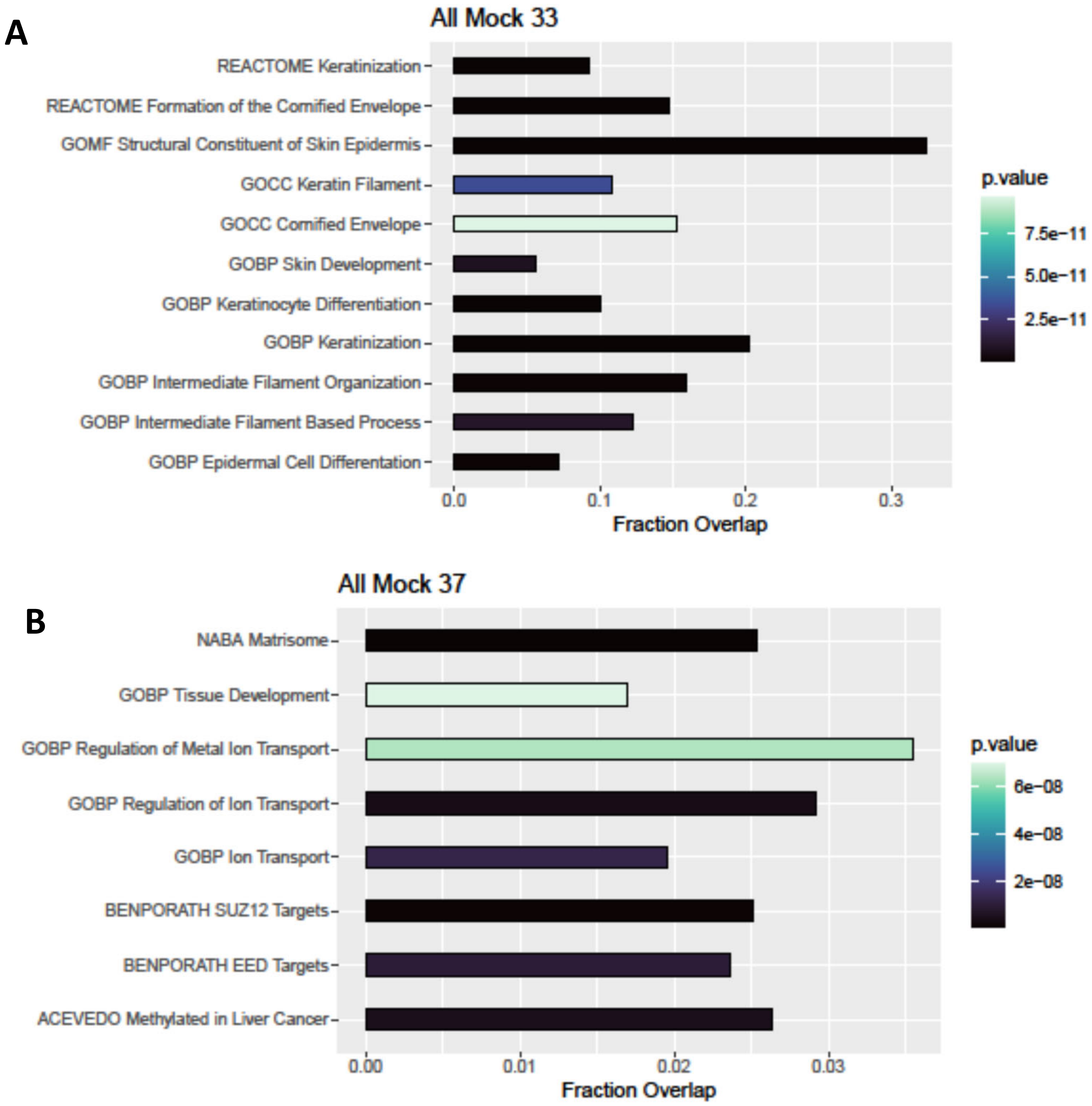


**Supp Fig 5.** *Biological Process Enrichment analysis for genes differentially expressed due to temperature*. Differential expression analysis was run between all (24 and 48HPI) mock infected samples to identify genes differentially regulated due to temperature. Biological Process enrichment analysis was run on the top 250 genes for 33$^{o}$C(A) or 37$^{o}$C (B). Fraction overlap indicates the number of the top 250 genes that were found to overlap with the indicated pathway divided by the total number of genes in that pathway.
