## Supplementary material for "Early transcriptional responses of human nasal epithelial cells to infection with Influenza A and SARS-CoV-2 virus differ and are influenced by physiological temperature": supp figure 6

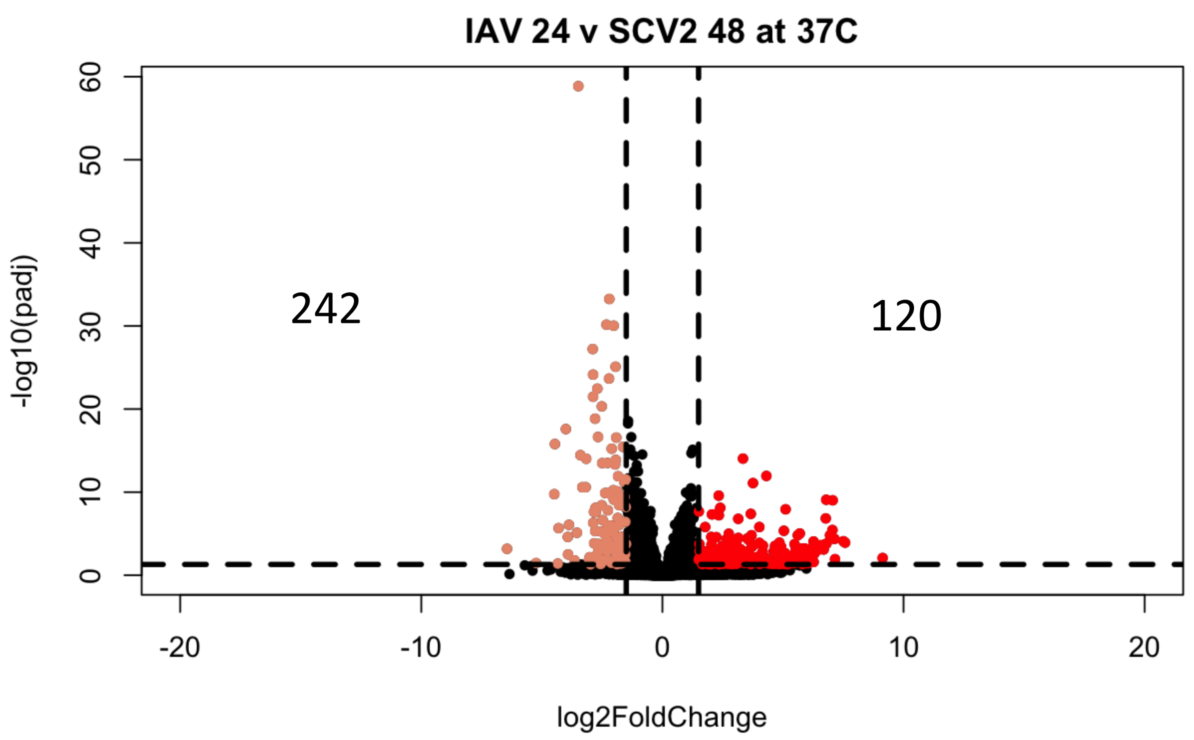


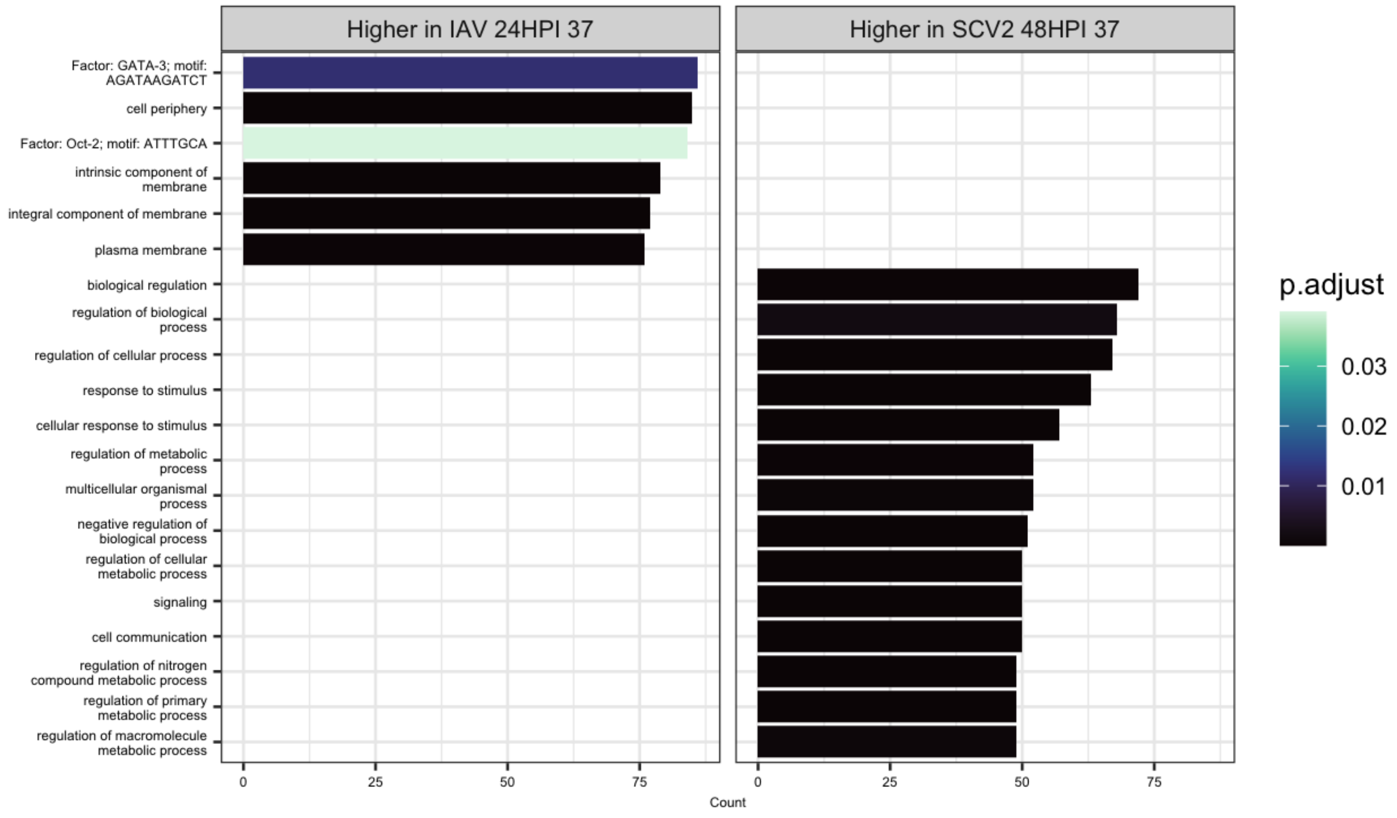


**Supp Fig 6.** *Test to see if transcriptomic changes due to virus used for infection are simply delayed or are qualitatively different*. Differential expression analysis was run between IAV 24HPI and SCV2 48HPI infected samples at 37ºC (A). Significantly differentially expressed genes ($p<0.05$, log2FC > 1.5) were then used in pathway enrichment analysis (B). The size of the bars indicate the number of genes in that pathway identified, and the bars are colored based on p value.
